# Laplacian Dynamics and Kron Reduction in Species-Reaction Graphs of Chemical Reaction Networks

**DOI:** 10.1101/2025.10.15.682662

**Authors:** Manvel Gasparyan, Upinder S. Bhalla, Ovidiu Radulescu, Shodhan Rao

## Abstract

We present a new method for deriving the dynamics of chemical reaction networks using the Laplacian matrix of the corresponding species–reaction graph, in contrast to previous works that use the Laplacian of the graph of complexes. Species-reaction graphs are bipartite graphs that contain two sets of vertices, one representing species and the other representing reactions, connected by directed edges that indicate relationships between them. Our approach starts by assigning appropriate edge weights to this bipartite graph, which are then used to compute the weighted graph Laplacian. This Laplacian reformulation of the system of differential equations governing the network dynamics emphasizes the flow of information throughout the chemical reaction network considered as causal network. As an application of this framework, we introduce a novel model reduction technique based on the Kron reduction of the weighted Laplacian matrix associated with the species-reaction graphs. Our systematic approach involves identifying nodes for deletion while preserving the bipartite structure, followed by constructing the Kron-reduced model. To demonstrate the effectiveness of our method, we apply it to a complex biochemical network, showing how model simplification facilitates analysis and interpretation of these systems.

## 1 Introduction

Chemical Reaction Networks (CRNs) serve as models for complex biochemical processes in cells and tissues. Studying these models is crucial for understanding diseases, developing new therapies, controlling bioengineering processes, and gaining insights into fundamental aspects of living systems [1, 2].

Models of CRNs are typically categorized as deterministic, stochastic, or hybrid. Deterministic models, commonly Ordinary Differential Equation (ODE) models [3] are valued for their simplicity, precision, and efficiency, particularly in systems with high reactant concentrations (large molecular counts, e.g., thousands per well-stirred volume), where random fluctuations in molecule numbers due to the inherent probabilistic nature of chemical reactions become insignificant compared to the average behavior. Stochastic models, such as Markov jump processes [4, 5], incorporate randomness to capture stochastic fluctuations in the species amounts. Though crucial for small-scale systems, these models are computationally intensive. Hybrid models [6, 7] combine deterministic and stochastic approaches, using deterministic methods for abundant species and stochastic methods for rare species, balancing accuracy and computational cost. This strategy reduces computational load while preserving the fluctuations of the species amounts. In this paper, we will focus solely on deterministic ODE models.

Graph theory offers powerful tools for analyzing CRNs. Network structures, such as (un)directed weighted graphs [8], multi-layer networks [9], temporal networks [10], and hierarchical graphs [11], facilitate the study of interactions in biochemical processes by highlighting structural properties such as feedback loops, key pathways, and their impact on qualitative dynamics, stability and robustness of the networks.

There is a growing need of mathematical tools for analyzing complex models of CRNs and for obtaining biological insight from the resulting models. The current state-of-the-art numerical tools for dynamical analysis of bioCRNs are known to suffer from a so-called curse-of-dimensionality. Therefore it is essential to develop mathematical techniques that can reduce a given kinetic model of a bioCRN to a simplified version that more or less mimics the behaviour of the original model, but has lesser number of parameters and differential equations.

For a comprehensive review of key model reduction strategies for CRNs, we refer the reader to [12] and [13]. Some of these model reduction methods have a graphical interpretation and can be described using graph rewriting operations, such as pruning (node elimination) and pooling (node aggregation). Among them, are the popular Quasi-Steady-State (QSS) and Quasi-Equilibrium (QE) approximations [14, 15], the exact lumping methods [16– 19] and the Kron reduction method [20–28]. In this paper, we revisit the latter in the context of a new formulation of CRN dynamics based on the Laplacian matrix of the Species-Reaction Graph (SR-graph).

Kron reduction [29] is a technique developed to simplify electrical grid models represented by directed graphs, by reducing network dimensionality while preserving voltage and current behavior at key nodes. It eliminates internal nodes, generating an equivalent network. The method was adapted for CRNs [20–28], simplifying CRNs represented by Graph of Complexess (GCs) (formal combinations of chemical species that appear together as reactants or products in a reaction) by eliminating some complexes. Following this adaptation, Kron reduction has also been employed as a tool for parameter estimation [23, 30] in CRN models.

The Kron reduction of the GC has limitations, as eliminating a complex may remove multiple species and reactions, potentially distorting the dynamics. For example, the CRN with the reactions *S*_1_ ⇄ *S*_2_ and *S*_2_+ *S*_3_ → *S*_4_ has four distinct complexes, namely { *S*_1_, *S*_2_, *S*_2_ + *S*_3_, *S*_4_}.

Eliminating the complex *S* _2_ + *S* _3_ would remove the second reaction entirely, leaving only the first one, which constitutes a large perturbation of the overall dynamics and is therefore not a good reduction. To address this, we propose a more flexible reduction using the SR-graph, a bipartite representation where species and reactions form disjoint node sets, with edges representing the relationships between them [31, 32].

This paper addresses two main problems: (i) establishing a link between the SR-graph and the system of ODEs derived from the stoichiometry and rate equations of a CRN, demonstrating how SR-graph edge weights based on stoichiometry and the state-dependent reaction rates can be used to reconstruct the ODE model; and (ii) proposing a novel model reduction technique that applies Kron reduction to the SR-graph of a CRN, rather than to the GC as done in prior work. Earlier, similar results were obtained for CRNs governed by Mass-Action Kinetics (MAK) [33]; here we generalize them to CRNs governed by general enzyme kinetics, which include MAK as a special case. The structure of the paper is as follows. In Section 2, we introduce the necessary techniques for deriving the main results of this work, including the standard framework for modeling CRNs – a system of ODEs based on stoichiometry. In Section 3, we present the Laplacian of a SR-graph and examine its relationship with the stoichiometry-based ODE model. In Section 4, we propose a novel model reduction technique that applies Kron reduction to the SR-graph of CRNs. Finally, in Section 5, we validate our reduction approach using a real-world CRN example. We apply it to an Activity-dependent Movement of a Glutamate Receptor (AMPAR) signaling network, used to model changes of synaptic strength in neuroscience [34]. Notably, our model reduction preserves the property of bistability which is a crucial property of this model. Our reduction method eliminates 35.71% of the species and 48% of the reactions in this model, while preserving accuracies of 1.73% and 0.34% for trajectories leading to each of the model’s two attractors.

**Notations:** ℝ^*n*^ and 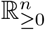 represents the *n*-dimensional Euclidean space and its non-negative orthant. For a vector **v** ∈ ℝ^*n*^, *v*_*i*_ ∈ ℝ refers to its *i*th entry, i.e., the vector **v** can be represented as 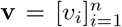. Vector of length *m* composed entirely of ones is denoted as **1**_*m*_. Additionally, **0**_*m*_ denotes the vector of length *m* all of whose entries are zeros, and **0**_*m*×*n*_ denotes the *m* × *n* matrix with each entry equal to zero. To avoid confusion, if there is no risk of ambiguity, we will use **0** in place of these notations. *A*_*ij*_ represents the entry of the matrix **A** that is located in the *i*th row and the *j*th column. For a finite set *I*, the number of elements in *I* is denoted by |*I*|, which represents the cardinality of the set. For a mutlivariable, real-valued function *φ* : ℝ^*n*^ → ℝ, *∂*_*i*_*φ* denotes the partial derivative of *φ* with respect to its *i*th variable.

## 2 Background

In this section, we follow the standard CRN framework to derive its mathematical formulation. We also introduce a technique for comparing the dynamics of the original and reduced CRN models, forming the basis of our modeling and reduction approach.

### 2.1 Stoichiometric mathematical models

We consider CRNs with only non-autocatalytic reactions; that is, no species appears simultaneously as both a substrate and a product of the same reaction. Autocatalysis is a rare phenomenon in biochemistry, and this phenomenon occurs only under special circumstances. Later on, in Section 4 of this manuscript, we will show that our reduction procedure leads to new reactions in which the complexes, i.e. the combinations of species of the left and right hand sides of the reactions of the original network are not retained in most cases. Hence our reduction procedure may lead to reactions that are not contained in the original CRN. In order to make the reactions of our reduced CRN acceptable to biochemists, we ensure that the reduced network arising from a given non-autocatalytic CRN in also non-autocatalytic.

Assume that *n*_*s*_ distinct chemical species *S*_*i*_, *i* = 1, …, *n*_*s*_, are involved in *n*_*r*_ irreversible reactions ℛ_*i*_, *i* = 1, …, *n*_*r*_. We do not exclude the possibility that reactions can be reversible. In this case, we consider each direction of a reversible reaction as a single separate reaction. Such a CRN can generally be written in the following form:

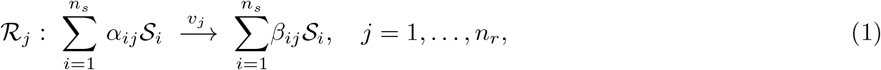

where *α*_*ij*_ and *β*_*ij*_ are non-negative constants representing the number of molecules of the *i*th species *S*_*i*_ in the substrates and the products of the *j*th reaction ℛ_*j*_. Here, *v*_*j*_ denotes the rate of the *j*th reaction ℛ_*j*_, which is a multivariable, real-valued, and non-negative function of the species’ concentration vector. Note that we do not exclude the possibility of having a reaction in which all the constants *α*_*ij*_ are zero simultaneously. For example, if there exists a reaction ℛ_*k*_ such that *α*_*ik*_ = 0 for all *i*, then the corresponding reaction is of the form 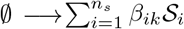, where ∅ indicates the absence of any reactant species. Such reactions represent inflows into the CRN, modeling the spontaneous creation or external supply of the product species involved. Similarly, it is also possible to have a reaction in which all the constants *β*_*ij*_ are zero. In this case, the reaction represents an outflow from the CRN and modeling the degradation or removal of the substrate species involved.

The connection between species and reactions is established through an *n*_*s*_ × *n*_*r*_ stoichiometric matrix, denoted by **S**. The elements of this matrix are defined as 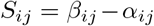 Let 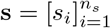 and 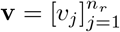 represent the vector of species concentrations and the vector of reaction rates, respectively. The fundamental framework characterizing the evolution of the species concentration vector is described by the following system of ODEs:

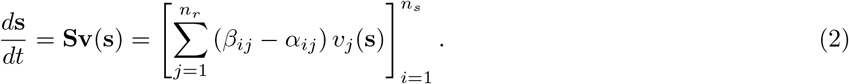

Buffered species—i.e., those that remain constant, do not participate as substrates or products, but act as modifiers—are not represented by an ODE of the form given in Eq. (2), as their concentration derivatives are zeros. However, they appear in the model through the rate expressions as constants, influencing reaction kinetics and potentially being treated as parameters. We further assume, that rates of zero order reactions are constant inputs to the system, i.e.

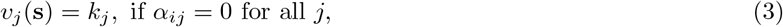

where *k*_*j*_ is a constant.

To illustrate the aforementioned approach for deriving the system of ODEs, we present a simple example of a CRN and demonstrate the modelling process.

#### Example 1.

*Consider the following example of a CRN of five distinct chemical species and three unidirectional reactions:*

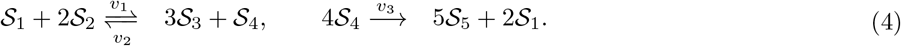

*The* 5 × 3 *stoichiometric matrix* **S** *of the CRN* (4) *is:*

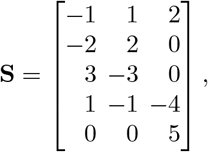

*and the corresponding system of ODEs can be expanded to:*

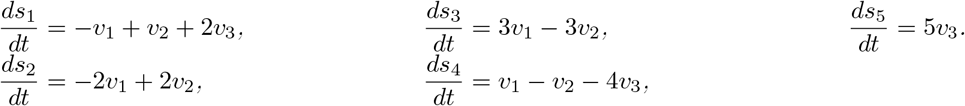

### 2.2 Enzyme kinetics law

Every CRN can be uniquely represented by a set of ODEs, as shown in Eq. (2), regardless of its rate governing laws, i.e. the governing principles for constructing the reaction rates as a function of the species concentrations. In this paper, we consider a set of rate governing laws known as General Enzyme Kinetics Law (GEKL) [20], that cover the most prevalent rate laws globally, governing numerous real-life bio-CRNs. According to this law, the rate *v*_*j*_ of the *j*th reaction is given by:

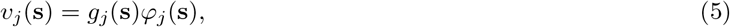

where *φ*_*j*_ is a multivariable monomial of the species concentration vector **s** defined by:

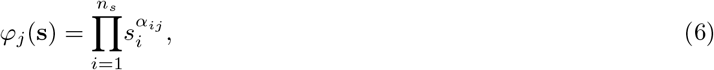

and *g*_*j*_ is a multivariable, real-valued, and non-negative rational function of its arguments.

MAK is an example of GEKL, where the rational terms are constant functions representing the rate constants of the corresponding reactions. A comprehensive list of well-known GEKLs, such as Michaelis-Menten kinetics [35] and Hill kinetics [36], can be found in the appendix of [37]. As a matter of fact, GEKL is very general, as it encompasses all kinetic laws that depend rationally on species concentrations.

Note that the reaction rate vector **v**, of a CRN governed by GEKL can be written as **v** = **Γ**_**g**_***φ***, where 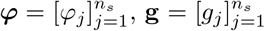, and **Γ**_**g**_ = diag(**g**). This representation is especially useful for improving computational efficiency and optimizing automation.

#### Example 2.

*With reference to the CRN depicted in Eq. (4), assuming its reactions are governed by GEKL, one possible formulation for the expressions of the reaction rates is as follows:*

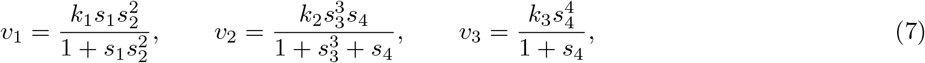

*where k*_*i*_ *are positive constants. In this case, the monomials corresponding to the reactions are given by:*

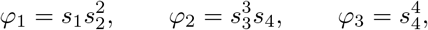

*and the rational terms within the expressions of the reaction rates are as follows:*

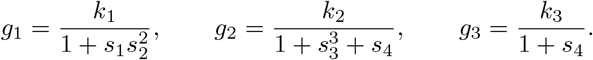

### 2.3 Species-reaction graphs

A SR-graph [31, 32] is a graphical framework for representing the dynamic relationships between species and reactions in a CRN. It provides insights into chemical transformations by illustrating the CRN structure.

The SR-graph of a CRN is a weighted directed bipartite graph (𝒱_*s*_, 𝒱_*r*_, ℰ), where 𝒱_*s*_ and 𝒱_*r*_ represent species and reaction nodes, respectively. Reversible reactions are modeled as two irreversible ones. The edges ℰ follow these rules: (i) if *S*_*i*_ is a reactant in reaction ℛ_*j*_, the edge goes from species *S*_*i*_ to reaction ℛ_*j*_; (ii) if *S*_*i*_ is a product of reaction ℛ_*j*_, the edge goes from reaction ℛ_*j*_ to species *S* _*i*_. Later, we will demonstrate how to appropriately assign the edge weights and explain their role in our model reduction procedure.

#### Example 3.

*To clarify the construction of the SR-graph for a given CRN, we use the example in Eq. (4).* The CRN consists of five species and three reactions, leading to a node set 𝒱 _*s*_ = { *S* _1_, …, *S* _5_} *for species and 𝒱* _*r*_ = { ℛ _1_, …, ℛ _3_} *for reactions. Since S* _1_ *is a reactant in* ℛ_1_, *there is a directed edge from S* _1_ *to* ℛ_1_ *in the SR-graph. Similarly, as S*_3_ *is a product in* ℛ_1_, *there is a directed edge from* ℛ_1_ *to S*_3_. *The remaining edges are constructed in the same manner. The resulting graph is shown in Fig. 1, where species nodes are represented as blue circles, reaction nodes as red squares, edges from species to reactions in orange, and edges from reactions to species in green*.

**Fig. 1:**
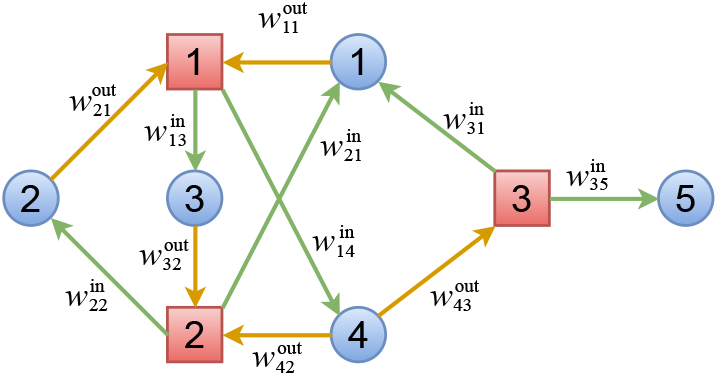
The SR-graph of the enzymatic CRN in Eq. (4), with species nodes (blue circles) and reaction nodes (red squares). Edges from species to reactions are orange, while those from reactions to species are green.

### 2.4 Symmetrized error integral

We present a metric designed to quantify the differences in the dynamical behavior between the original model and its corresponding reduced model. To achieve this, we utilize the error integral.

#### Definition 1

(Error integral). *Let* 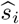 *and s*_*i*_ *represent the concentrations of the ith species in the reduced model and the original model, respectively. Further let T*_*i*_ *denote the duration of the time interval during which we examine the differences in the concentration behavior of the ith species between a base trajectory of the original model and the corresponding trajectory of the reduced model. The error integral between the original model and the reduced model is defined as:*

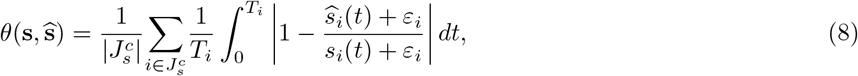

*where* 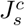 *is the set of indices in the original model corresponding to the species that are preserved in the reduced model and ε*_*i*_ *is a small positive value used to prevent division by zero and large errors that may arise when* 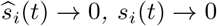.

The concept of the error integral is adapted from [20], where all *T*_*i*_ are assigned the same value. In contrast, this study allows each species to be evaluated over its own time interval. For zero-deficiency CRNs, a suitable choice for *T*_*i*_ is the settling time of the *i*th species—defined as the time it takes for its concentration to enter and remain within a small, predefined neighborhood of its steady state 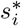. An automated method for computing these settling times is presented in [22, 23], where *T*_*i*_ is determined as the earliest time the concentration *s*_*i*_ enters this neighborhood and stays within it for the remainder of the simulation. The overall network settling time is then given by max_*i*_ *T*_*i*_, which is used uniformly for all species in [20].

It is important to note that the error integral, as defined in Eq. (8), is not symmetric. This lack of symmetry arises because the measure is dependent on the direction of comparison between the full and reduced models; the error quantified when comparing the full model to the reduced one is not necessarily the same as when the comparison is reversed. A symmetric error integral, on the other hand, is advantageous because it treats the discrepancies between the full and reduced models equally, regardless of the direction of comparison. For this purpose, we use the symmetrized version of the error integral, which was first introduced and applied in [38].

#### Definition 2

(Symmetrized error integral). *Let θ be the error integral between the original model and the reduced model, as defined in Eq. (8). The symmetrized error integral between the original model and the reduced model is defined as:*

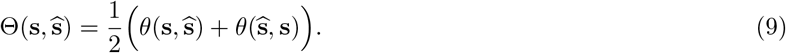

This symmetry provides a more balanced and comprehensive assessment of the differences in dynamical behavior between the two models, ensuring that both overestimations and underestimations are accounted for equivalently.

### 2.5 Schur complement

The Schur complement [39] plays a central role in Kron reduction, as it naturally arises from the Gaussian elimination of variables in linear systems of equations. To illustrate this, let us consider a block-partitioned linear system and show how one set of variables can be expressed in terms of the others, leading to the Schur complement. Consider the linear system:

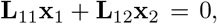

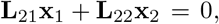

where **L**_11_ ∈ ℝ^*n*×*n*^, **L**_12_ ∈ ℝ^*n*×*m*^, **L**_21_ ∈ ℝ^*m*×*n*^, and **L**_22_ ∈ ℝ^*m*×*m*^. Assuming that **L**_22_ is invertible, we can express **x**_2_ in terms of **x**_1_ as 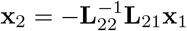. Substituting this relation into Eq. (10) yields the reduced equation:

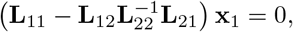

which illustrates the construction of the Schur complement.

#### Definition 3

(Schur complement). *Let* **L**_11_ ∈ ℝ^*n*×*n*^, **L**_12_ ∈ ℝ^*n*×*m*^, **L**_21_ ∈ ℝ^*m*×*n*^, *and* **L**_22_ ∈ ℝ^*m*×*m*^ *be constant matrices such that* **L**_22_ *is invertible. Consider the following* (*n* + *m*) × (*n* + *m*) *block matrix:*

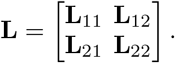

*The Schur complement of the block* **L**_22_ *of the matrix* **L** *is the n* × *n matrix* 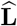 *defined as:*

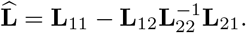

#### Remark 1.

*The definition of the Schur complement requires the block* **L**_22_ *to be invertible. If* **L**_22_ *is singular, the classical Schur complement is not defined. In such cases, a common generalization replaces the inverse* 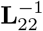 *with the Moore–Penrose pseudoinverse* 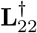, *i*.*e* 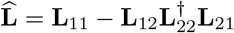, *leading to the generalized Schur complement*.

## 3 Dynamics on a species-reaction graph

In this section we present one of our study’s key findings: we show how to derive a new representation of the enzymatic CRNs using their SR-graphs. By defining edge weights, we utilize the graph Laplacian of the SR-graph to construct a system of ODEs that accurately reflect the CRN dynamics.

### 3.1 Graph Laplacian

We now demonstrate how to appropriately define the weights of the edges in the SR-graph and subsequently explain the construction of the weighted directed Laplacian matrix of the SR-graph associated with the given CRN using the weighted directed adjacency matrix. We define the weights of the edges as follows:

1. For an edge with the node corresponding to species *S*_*i*_ as the source and the node corresponding to reaction ℛ_*j*_ as the target, the weight of this edge is 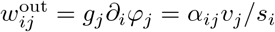. Here, as previously described, *g*_*j*_ and *φ*_*j*_ denote the rational and the monomial terms, respectively, in the expression for the rate *v*_*j*_ of reaction ℛ_*j*_.
2. For an edge with the node corresponding to reaction ℛ_*j*_ as the source and the node corresponding to species *S*_*i*_ as the target, the weight 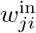 of this edge is the number of molecules of species *S*_*i*_ resulting from the reaction ℛ_*j*_, namely 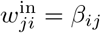.

Let us now re-index the *n*_*s*_ species and *n*_*r*_ reactions as nodes of the SR-graph using the following substitution: *i* → *i*, for *i* ∈ *S*, and *j* → *j* + *n*_*s*_, for *j* ∈ ℛ. In the new indexation, the species use the indices 1, …, *n*_*s*_ and reactions the indices *n*_*s*_ + 1, …, *n*_*s*_ + *n*_*r*_.

The adjacency matrix **A**(**s**) of the SR-graph associated with a CRN is a (*n*_*s*_ + *n*_*r*_) × (*n*_*s*_ + *n*_*r*_) matrix, such that its entry *A*_*mk*_ is equal to the weight of the edge having the *k*th node as the source node and *m*th node as the target node. In the case when there is no such edge, we have *A*_*mk*_ = 0. Note that the definition of the adjacency matrix we use is the transpose of the adjacency matrix typically defined in the literature. To minimize confusion and avoid excessive notation, we will adopt our definition throughout, without any loss of generality. More precisely, one has:

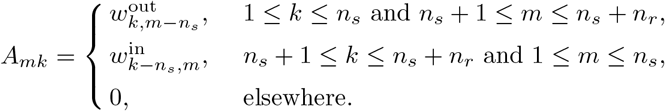

The weighted directed degree matrix **D**(**s**) of the SR-graph is a (*n*_*s*_ + *n*_*r*_) × (*n*_*s*_ + *n*_*r*_) diagonal matrix such that its *m*th diagonal entry is equal to the sum of the elements of the *m*th column of the weighted adjacency matrix **A**(**s**), namely:

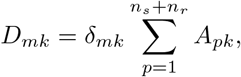

where *δ*_*mk*_ is the Kronecker delta.

The Laplacian matrix is an (*n*_*s*_ + *n*_*r*_) × (*n*_*s*_ + *n*_*r*_) square matrix defined as:

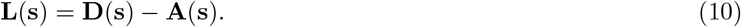

The Laplacian matrix can be described as follows: for distinct nodes *m ≠ k*, if there is an edge from the *k*th node to the *m*th node, *L*_*mk*_ is the negative of the weight of that edge. Otherwise, *L*_*mk*_ = 0. For the diagonal entries *m* = *k*, we set *L*_*kk*_ = *−* ∑_*m ≠k*_ *L*_*mk*_, such that the column-wise sum of **L**(**s**) is the zero vector. Note that the main *n*_*s*_ × *n*_*s*_ block matrix of **L**(**s**) is a diagonal matrix. This is because in the SR-graph there are no edges connecting two species nodes. Similarly, the right-lower *n*_*r*_ × *n*_*r*_ block matrix is also diagonal because there are no edges connecting two reaction nodes. The other entries of the Laplacian matrix can be described as follows. If there is a reaction ℛ_*j*_ for which the species *S*_*i*_ is a substrate, then the off-diagonal element 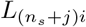 is equal to the negative product of the rational term in the rate expression for the *j*th reaction ℛ_*j*_ and the partial derivative of the corresponding monomial with respect to the concentration of the *i*th species *S* _*i*_, i.e. 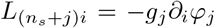. If there is a reaction ℛ _*j*_ for which the *i*th species *S* _*i*_ is a product, then the off diagonal element 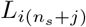 is the number of molecules of the *i*th species *S* _*i*_ in the expression of the *j*th reaction ℛ _*j*_, i.e. 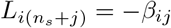.

Any directed graph is defined by an incidence matrix, which represents the connections between its vertices and edges. In the case of a SR-graph, the (*n*_*s*_ + *n*_*r*_) × *n*_*e*_ incidence matrix **B**, with *n*_*e*_ being the number of edges, is defined as follows:

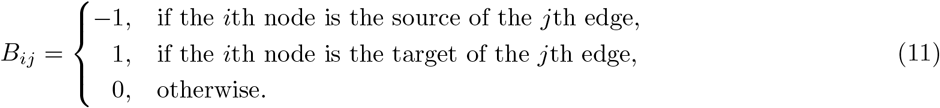

Define the (*n*_*s*_ + *n*_*r*_) × *n*_*e*_ outgoing matrix **Δ** of the SR-graph by:

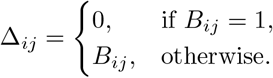

It can be shown that the Laplacian matrix is given in terms of matrix multiplication by:

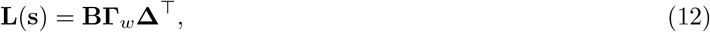

where **Γ**_**w**_ = diag(**w**) is the conductance matrix with **w** being the vector of weights.

#### Example 4.

*Consider again the CRN example presented in Eq. (4) and its corresponding SR-graph visualized in Fig. 1. By assigning weights to the edges according to the prescribed rule, the resulting edge weights are as follows:*

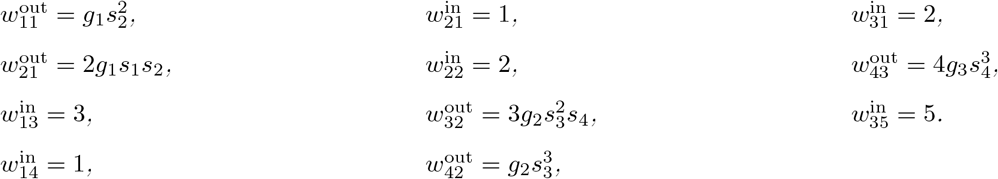

*The weighted adjacency matrix* **A**(**s**) *and the weighted degree matrix* **D**(**s**), *both of size* 8 × 8, *are as follows:*

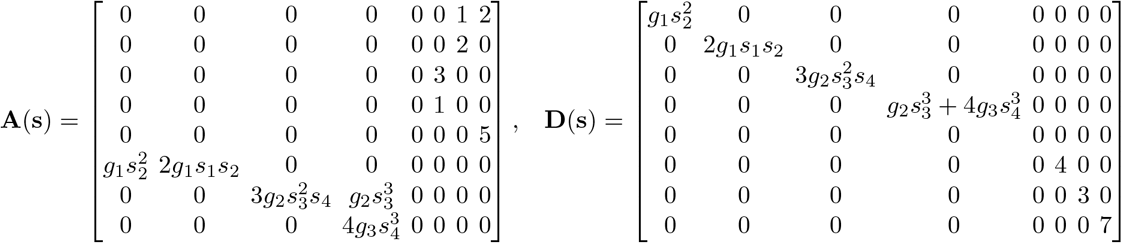

*and the* 8 × 8 *weighted directed Laplacian matrix* **L**(**s**) *is:*

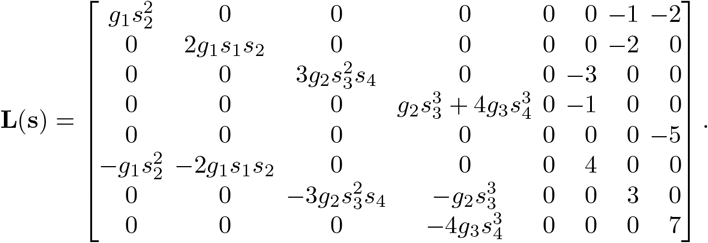

*Note that the Laplacian matrix can also be represented in the matrix multiplication form given in Eq. (12) with the* 8 × 11 *incidence matrix* **B** *and the* 8 × 11 *outgoing matrix* **Δ** *given as follows:*

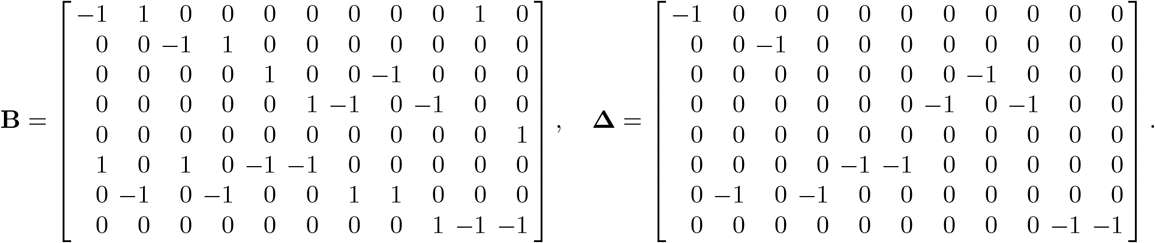

### 3.2 Properties of Laplacian matrices of SR-graphs

Beyond the standard properties common to all Laplacian matrices, the Laplacians of SR-graphs constructed using our weight assignment scheme exhibit additional structural features. In particular, certain block structures within these matrices take on specific forms, owing to the bipartite nature of the SR-graphs and the characteristics of the assigned weights. We summarize these additional properties in the following. The (*n*_*s*_ + *n*_*r*_) × (*n*_*s*_ + *n*_*r*_) Laplacian matrix has the following block structure:

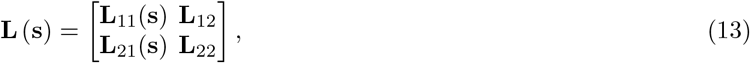

where:

*P1)* **L** (**s**) possesses the standard properties of Laplacian matrices, that is, for every 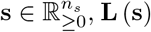 has non-negative diagonal entries, non-positive off-diagonal entries, and zero column sums.

*P2)* **L**_11_(**s**) is an *n*_*s*_ × *n*_*s*_ diagonal matrix with entries depending on the vector of species concentrations **s**. The matrix is diagonal because, due to the bipartite structure of the SR-graph, there are no edges connecting species nodes to one another.

*P3)* **L**_12_ is an *n*_*s*_ × *n*_*r*_ constant matrix with integer entries. This is due to the fact that the weights of the edges from reaction nodes to species nodes in the SR-graph are constants. Specifically, [**L**_12_]_*ij*_ = *−β*_*ij*_, where *β*_*ij*_ is the stoichiometric coefficient of the *i*th species in the product of the *j*th reaction.

*P4)* **L**_22_ is an *n*_*r*_ × *n*_*r*_ constant diagonal matrix. The matrix is diagonal because, again, the bipartite structure of the SR-graph prevents any edges between reaction nodes. Specifically 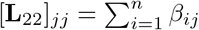

*P5)* **L**_21_(**s**) is an *n*_*r*_ × *n*_*s*_ matrix with entries depending on **s**. For each row *j* of **L**_21_(**s**), there is a rational function *g*_j_ (**s**) and a monomial 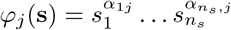 such that

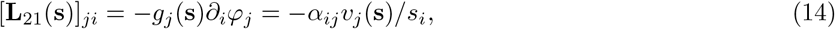

where *v*_*j*_(**s**) = *g*_*j*_(**s**)*φ*_*j*_(**s**). This results from (5). In other words, each row of **L**_21_(**s**) is the product of a rational integrating factor and the gradient of a monomial with respect to **s**.

*P6)* In the Laplacian matrix **L**(**s**), there is no *j* ∈ {1, …, *n*_*s*_ + *n*_*r*_} such that the *j*th row and the *j*th column are entirely filled with zeros. This is because we consider CRNs with connected SR-graphs.

*P7)* Since we consider CRNs with non-autocatalytic reactions, the SR-graphs cannot contain both an edge from a species to a reaction and the reverse edge simultaneously. Consequently, the entries of the Laplacian matrix satisfy the following property: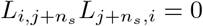, for all *i* = 1, …, *n*_*s*_ and *j* = 1, …, *n*_*r*_.

#### Remark 2.

*Note that, a matrix is the Laplacian matrix of the SR-graph of a CRN if it satisfies properties P1– P5 listed above. Also note that a CRN with non-integer rational stoichiometric coefficients may be modified so as to satisfy condition* P3 *without altering its dynamics. Assuming that the j*^*th*^ *reaction has non-integer rational coefficients, let q denote the least common multiple of the denominators of the coefficients α*_*ij*_ *and β*_*ij*_ *for i* = 1, …, *n*_*s*_. *The transformation α*_*ij*_ ↦ *qα*_*ij*_, *β*_*ij*_ *↦ qβ*_*ij*_ *and* 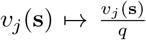 *produces an equivalent reaction (i*.*e*., *a reaction with the same dynamics) with integer stoichiometric coefficients (see [23, Lemma 1]). The properties P6 and P7 have been included because for our reduction, we only consider autocatalytic CRNs with connected SR-graphs*.

### 3.3 Laplacian-based dynamical model

Here, we demonstrate how to derive the dynamical system given in Eq. (2) and the corresponding enzymatic reaction rates from the SR-graphs by utilizing its Laplacian matrix. We first introduce the vector of *net stoichiometric imbalances* of reactions. It is a vector 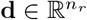, where the net stoichiometric imbalance of the reaction 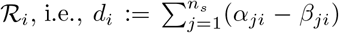, represents the difference between the total number of molecules consumed and the total number produced in the reaction ℛ _*i*_. Moreover, *d*_*i*_*v*_*i*_ represents the net rate of change of the total species concentration due to the reaction ℛ _*i*_. Therefore, reactions with zero net stoichiometric imbalance conserve the total species concentration.

#### Theorem 1.

*Let* 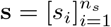 *and* 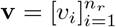 *denote the vector of species concentrations and the vector of reaction rates. Further let* **Γ**_**d**_ = diag(**d**), *with* 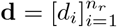 *being the vector of net stoichiometric imbalances of the reactions*.

*Then the following identity holds:*

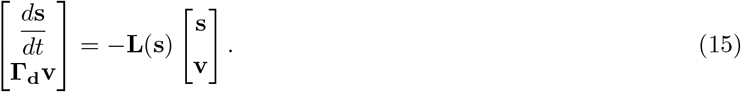

*Proof*. Let **Λ** denote the right-hand side of Eq. (15). For *i* = 1, …, *n*_*s*_, the *i*th component of **Λ** can be rewritten as:

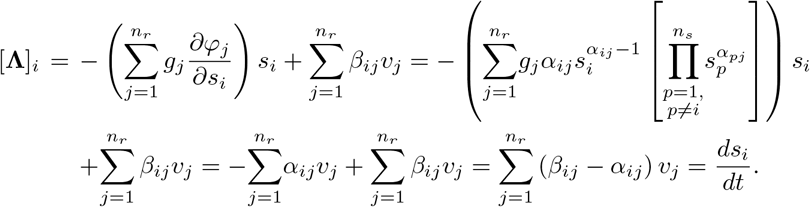

The last identity follows form Eq. (2). Concerning the remaining components of **Λ**, for *j* = 1, …, *n*_*r*_, we have:

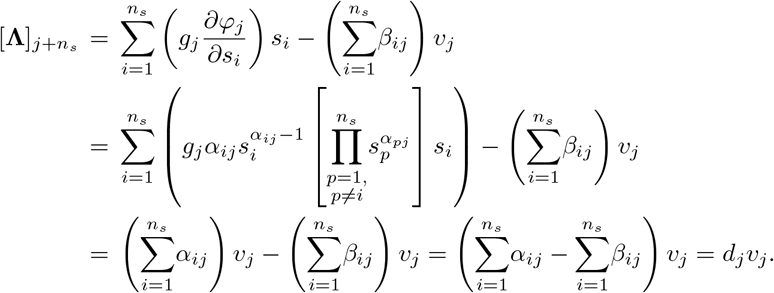

This concludes the proof.

Given a model of the form presented in Eq. (15), we can derive the expressions for the reaction rates **v** as functions of the species concentrations **s**, as well as the corresponding system of ODEs provided in Eq. (2), as detailed in the following remark.

#### Remark 3.

*Consider the block decomposition of the Laplacian matrix* **L**(**s**) *given in EQ. (13). The key argument is that the last equations of Eq. (15), which correspond to* **v**, *depend linearly on* **v**, *allowing* **v** *to be determined using Gaussian elimination. Consequently, Eq. (15) implies that:*

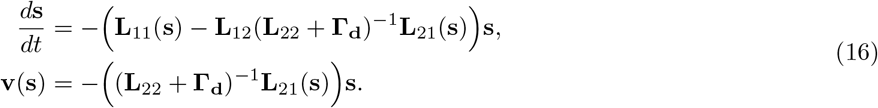

*Therefore, given the expression on the right-hand side of Eq. (15), simple algebraic manipulations can be employed to derive both the reaction rate vector and the system of ODEs outlined in Eq. (2). One can recognize in the expression of* 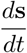 *the Schur complement of:*

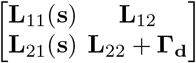

*corresponding to the block* **L**_22_ +**Γ**_**d**_. *Note that the matrix* **L**_22_ +**Γ**_**d**_ *is diagonal, i*.*e* 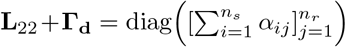. *This implies that all reaction rates can be computed using equation* (16), *except for zero-order reactions, which involve no reactants. However, this has no impact if the rates of zero-order reactions are treated as input parameters, see Eq. (3)*.

This novel Laplacian-based and bipartite graph representation of CRNs provides a structured framework for analyzing species-reaction interactions. By explicitly capturing the network’s topology, it offers valuable insights into both its structural and dynamic properties. The Laplacian matrix governs the diffusion of information in complex networks, with its eigenvalues and eigenvectors determining how perturbations spread and at what rate [40, 41]. In the context of CRNs, this corresponds to the propagation of concentration changes through reaction-mediated interactions [42]. This perspective can inform strategies for assessing system stability, identifying key pathways, and optimizing network behavior, with applications in synthetic biology, systems biology, and biochemical engineering. While further theoretical developments are left for future work, we demonstrate the practical utility of this approach for model reduction.

## 4 Model reduction

A real-life CRN model is often complex, involving many variables and parameters that require substantial computational resources for analysis [43]. Model reduction addresses this by simplifying the model, retaining key dynamical features while reducing the number of variables and parameters for easier analysis.

We apply our novel modeling approach to reduce CRNs, using the SR-graph and its Laplacian matrix to simplify the system of ODEs. Our reduction method for CRNs governed by GEKL, draws inspiration from the Kron reduction technique for resistive electrical networks [29], and is supported by insights from related CRN studies [20–22].

### 4.1 Methodology Outline

The goal of our model reduction is to remove a specific set of species and reactions from the SR-graph, ensuring that the resulting directed graph remains a valid SR-graph. Removing a reaction node *j* is equivalent to enforcing that *d*_*j*_*v*_*j*_ = 0. The application of these balance constraints lead to a reduced model. Because the constraints are applied formally, their (approximate) validity must be tested. This is done numerically using the symmetrized error integral introduced in Section 2.4. We also require that the reduced model retains a SR-graph structure. To ensure this, we verify symbolically that the Schur complement Laplacian satisfies the properties listed in Section 3.2, as outlined in Algorithm 2.

Consider a CRN with dynamics given by Eq. (15). Let *I*_*s*_ and *I*_*r*_ represent the set of species and reaction indices, respectively. Further, let *J*_*s*_ and *J*_*r*_ denote the sets of indices of species and reactions to be deleted from the SR-graph, with 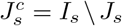 and 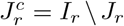. Define the following partitions for the concentration and reaction rate vectors.

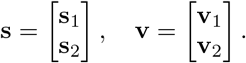

Here, **s**_1_ is the vector of length 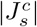 representing the concentrations of the species remaining in the model, and **s**_2_ is the vector of length |*J*_*s*_| representing the concentrations of the species being removed from the model. Similarly, **v**_1_ is the vector of length 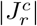 representing the rates of the reactions remaining in the model, and **v**_2_ is the vector of length |*J*_*r*_| representing the rates of the reactions being removed from the model. In this case, the model given in Eq. (15) can be rewritten as follows:

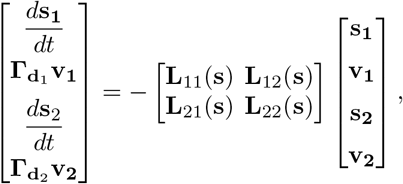

where **d**_*i*_ is the vector of stoichiometric imbalances corresponding to the reactions with rates given by **v**_*i*_. Here, the blocks **L**_11_(**s**), **L**_12_(**s**), **L**_21_(**s**), and **L**_22_(**s**) of the Laplacian matrix **L**(**s**) are matrices of sizes 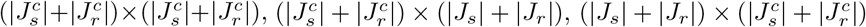, and (|*J*_*s*_| + |*J*_*r*_|) × (|*J*_*s*_| + |*J*_*r*_|), respectively.

Our model reduction procedure is based on the following heuristics. Deleting the species nodes *J*_*s*_ is equivalent to assuming that the concentration vector **s**_2_ satisfy the equation 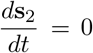. We therefore assume that the net effect of all the reactions on the species that have been removed is balanced, i.e., that the removed species are in a quasi-steady state. Similarly, deleting reaction nodes *J*_*r*_ is equivalent to assuming that the eliminated reactions **v**_2_ satisfy the equation 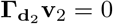. Our model reduction technique has similarities with both the QSS and the QE approaches, which are elaborated in the Supplementary Material.

The reduced model reads:

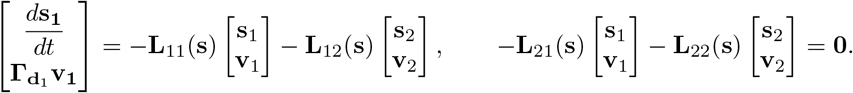

From the second equation we find that:

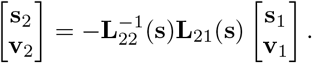

Substituting this result into the first equation yields:

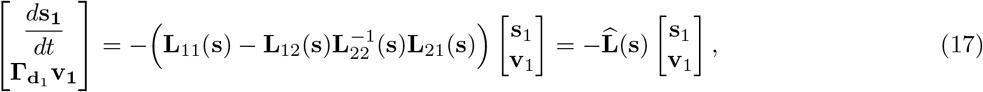

where 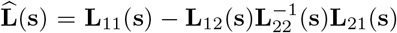. Note that 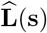 is the Schur complement of **L**(**s**) with respect to the indices corresponding to *J*_*s*_ ∪ *J*_*r*_. The reduction procedure does not apply when the matrix **L**_22_(**s**) is singular.

It is important to note that the reduced model remains independent of the order in which species and reactions are deleted. The matrix 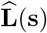 corresponds to a SR-graph where the species nodes correspond to the species with indices 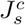 and the reaction nodes correspond to the reactions with indices 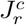. The theorem that follows establishes that 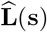 adheres to the first property mentioned in Section 3.2, expected from a weighted Laplacian matrix within the context of a SR-graph.

#### Theorem 2.

*Let* 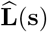 *be the Laplacian matrix of the SR-graph associated with the CRN. Let I* ⊂ {1, …, *n*_*s*_ + *n*_*r*_} *and* **L**(**s**) *be the Schur complement of* **L**(**s**) *with respect to the set of indices I. Then* **L**(**s**) *is again a Laplacian matrix, i*.*e*., *the following properties hold:*

1. *All non-diagonal entries of* 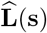*are non-positive for* 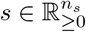.
2. *All diagonal entries of* 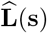 *are non-negative for*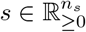.
3. 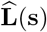*has zero column sums*.

*Proof*. The proof of the theorem directly follows from the proof provided in [44].

#### Remark 4.

*Theorem 2 establishes that the Schur complement of the Laplacian matrix is again a Laplacian matrix of an ordinary graph, implying that it need not correspond to a bipartite SR-graph. In the reduction algorithm described in the next section, we therefore additionally verify that the Schur complement satisfies the structural properties listed in Section 3.2, which ensure that the reduced matrix corresponds to a valid connected SR-graph with non-autocatalytic reactions*.

#### Remark 5.

*In Eq. (17), the Schur complement eliminates the reaction rates* **v**_2_, *but the remaining Laplacian blocks, and thus* 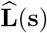, *still depend on both* **s**_1_ *and* **s**_2_. *To reduce the model by eliminating* **s**_2_, *one must solve the quasi-stationarity equation:*

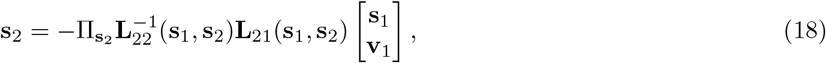

*where* 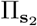 *projects onto* **s**_2_. *Substituting this into* 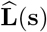, *the reaction rates* **v**_1_ *are obtained from Eq. (17) and expressed in terms of* **s**_1_ *via Gaussian elimination, as in Remark 3*.

*First note that, in some cases, Eq. (18) can be solved analytically to give an explicit expression for* **s**_2_ *in terms of* **s**_1_ *and* **v**_1_. *In such cases, we can obtain a reduced model in an analytical form. This corresponds to the cases where the vector of reaction rates* **v**_1_ *of the reduced model can be expressed as an analytical function of* **s**_1_. *However, in other cases where Eq. (18) cannot be analytically solved, it still can be solved numerically. In such cases, we cannot obtain analytical expressions for vector of the reaction rates* **v**_1_ *of the reduced model in terms of* **s**_1_. *The reduced model will be a computational model which is capable of producing the output trajectories for given initial conditions of species concentrations. It is important to note that other model reduction methods involving species elimination, such as the QSS and QE approximations, face similar challenges*.

#### Algorithm 1

CandidateCombinations

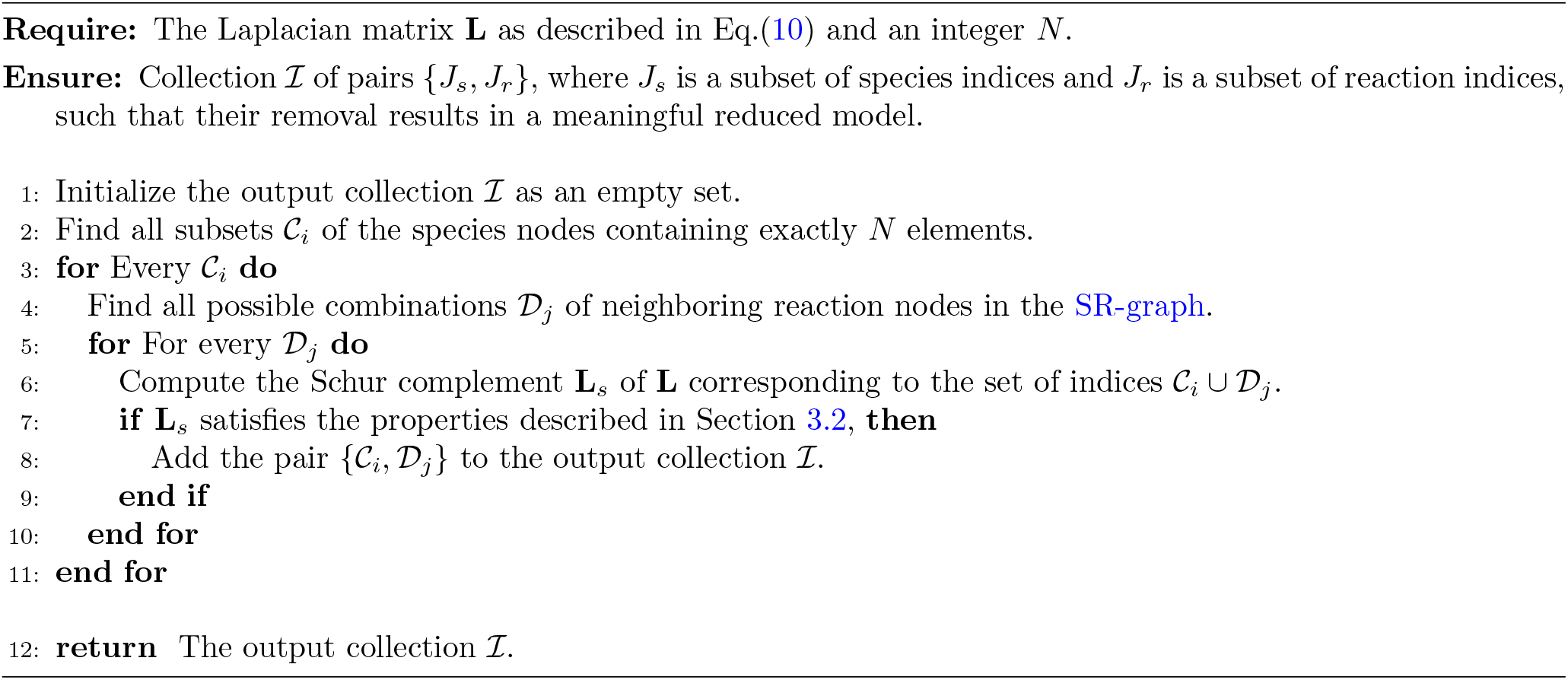

### 4.2 Reduction algorithm

In this section, we present the reduction algorithm, outlining the key steps and principles. We demonstrate how to identify the optimal subset of species and reactions for removal, ensuring that the resulting bipartite graph remains a valid SR-graph. A step-by-step reduction procedure applied to a simple CRN example is provided in the supplementary material.

#### Combination of candidate species and reactions for effective removal

We now outline the methodology for selecting candidate species and reactions for removal from the SR-graph via Kron reduction.

The core idea is to select a combination of species and reaction subsets such that the corresponding Schur complement of the Laplacian, associated with these subsets, remains a valid reduced Laplacian matrix – meaning that it satisfies the conditions stated in Section 3.2. Ideally, we would examine all possible combinations of species and reactions. However, this is computationally expensive. With *n*_*s*_ species and *n*_*r*_ reactions, the number of combinations would be 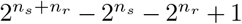, which is prohibitive for large values of *n*_*s*_ and *n*_*r*_. Thus, we need a more feasible approach to identify candidate subsets for removal.

To implement this procedure, we begin by selecting an integer *N*, which specifies the number of species nodes in each candidate subset. We then enumerate all distinct subsets 𝒞_*i*_ of species nodes containing exactly *N* elements. For each subset 𝒞_*i*_, we identify all possible combinations 𝒟_*j*_ of neighboring reaction nodes. A subset 𝒞_*i*_ ∪ 𝒟_*j*_ is considered a candidate for removal if its elimination results in a Schur complement satisfying the criteria of a valid Laplacian matrix, as presented in the Algorithm 1. This process is repeated for increasing values of *N*, starting with *N* = 1, systematically identifying increasingly larger candidate subsets. Since the symmetrized error integral tends to increase with *N*, the procedure is terminated when the error exceeds a predefined threshold.

#### Main reduction algorithm

We now present the main reduction algorithm, which is also schematized in Algorithm 2. Consider a given enzymatic CRN described by its stoichiometric matrix 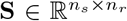 and the vector of reaction rates **v** as a function of species concentrations. The first step is to determine the SR-graph, which can be derived from the stoichiometric matrix as follows: the species nodes correspond to the rows, and the reaction nodes correspond to the columns of **S**. The edges are constructed based on the entries of *S*_*ij*_, where *i* is the row index and *j* is the column index. If the entry *S*_*ij*_ is negative, there is a directed edge from the *i*th species node to the *j*th reaction node in the SR-graph. If *S*_*ij*_ is positive, there is a directed edge from the *j*th reaction node to the *i*th species node. The weights of these edges are determined from the stoichiometric matrix and the vector of reaction rates, as explained at the beginning of Section 3.1.

Starting with the original model, we first generate candidate subsets of species and reactions for removal, as outlined in Algorithm 1. Among these, we select the subset whose deletion minimizes the value of the symmetrized error integral given in Eq. (9). We then apply Kron reduction to obtain the reduced model. This procedure is repeated iteratively on the reduced model: in each iteration, candidate subsets are determined, ranked, and the subset yielding the smallest symmetrized error integral is removed. The iteration is stopped when the symmetrized error integral of the reduced model (compared to the initial model) obtained at the end of an iteration exceeds a certain cutoff threshold. The reduced model obtained at the end of the previous iteration is then considered the final reduced model.

##### Algorithm 2

KronReduction

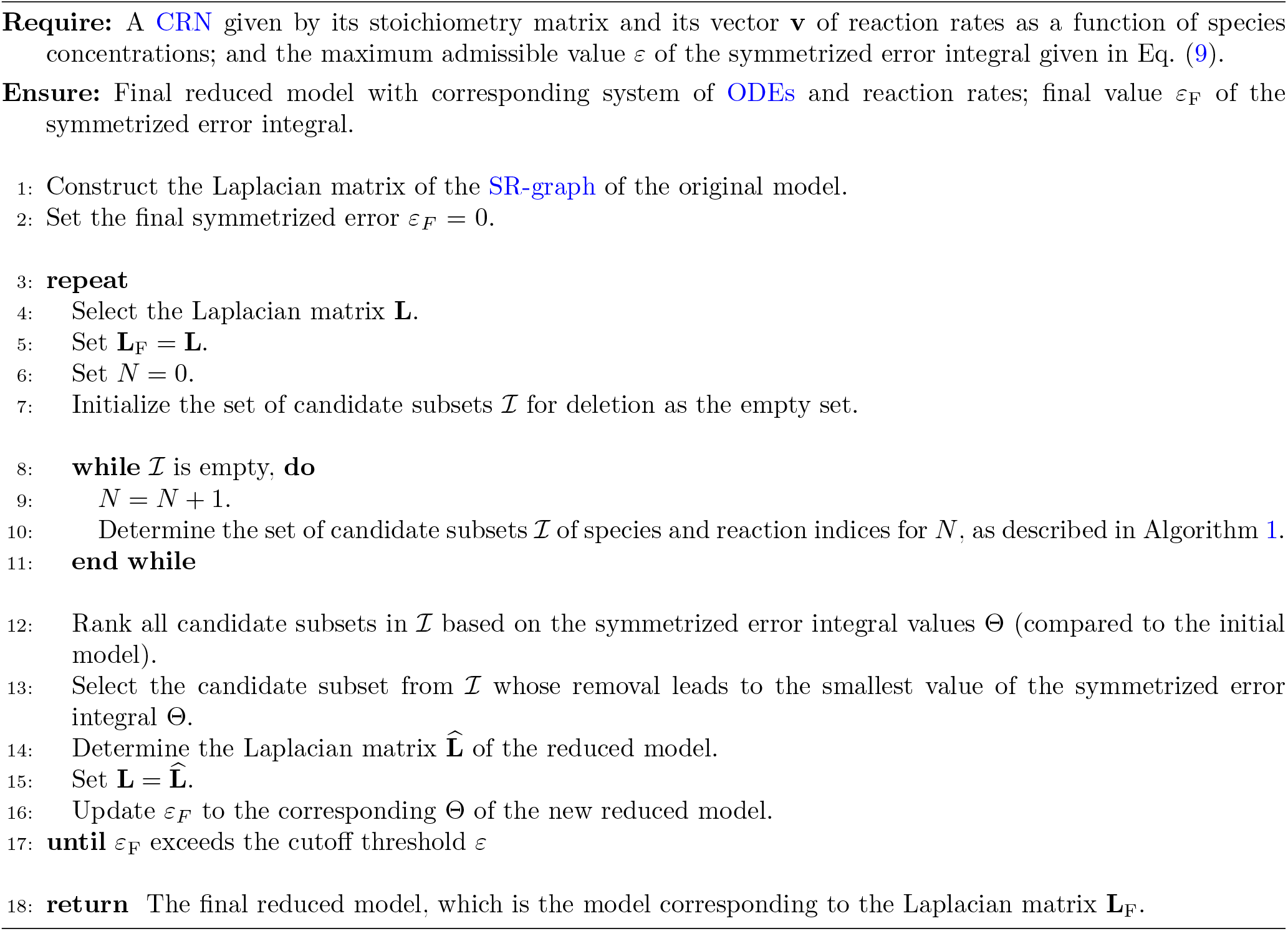

#### Automated Model Reduction Tools

We have developed Python and MATLAB packages for the automatic reduction of CRNs. These packages are provided as supplementary material and are publicly accessible via the following GitHub repository: https://github.com/manvelgasparyan/LK red. The tools require the biochemical model information, including the stoichiometric matrix and the expressions of the rate functions with available parameter values. In addition, a user-defined parameter must be specified, which is the maximum admissible threshold for the symmetrized error integral (15% in our examples, as detailed in the next section).

After submission, the tool processes the model and produces several outputs, including a list of removed species and reactions, the symmetrized error integral – which quantifies the discrepancy between the original and reduced models – the SR-graphs of both the models, and a visual comparison of their base trajectories. This suite of features ensures efficient model reduction while preserving key dynamical properties within a pre-specified tolerance.

## 5 Application of model reduction to a real-life example

In this section, we demonstrate the practical utility of our modeling and model reduction methods through a real-life CRN example. These techniques simplify complex biological systems while preserving model accuracy. The case study showcases how our approach can make computational models more manageable without losing essential dynamics, validating the applicability of our methods for analyzing and optimizing biological systems. We examine the AMPAR trafficking cycle model, originally derived from [34]. In addition, three further real-world examples, namely a model of N-(1-deoxy-D-fructose-1-yl)-glycine (DFG) thermal decomposition [45], a model of Hedgehog Signaling Pathway (HSP) [46], and a model of Cell Division Cycle (CDC) [47], are provided in the Supplementary Material. These models are available from the BioModels database at https://www.ebi.ac.uk/biomodels.

The considered AMPAR trafficking cycle model forms a bistable system in which the low state has few (∼ 10) membrane bound AMPA receptors (MR, MR* and MR** in the equations below), and the high state has large numbers (∼164 receptors). A brief increase in Protein Kinase A (PKA) promotes the model to the high state, representing synaptic potentiation, and a temporary reduction of PKA to near zero pulls the model from high to low state.

The original model comprises 2 buffered species and 14 non-buffered species. We decompose each reversible reaction into two unidirectional reactions, resulting in a total of 25 unidirectional reactions. A schematic representation of the CRN is shown in Fig. 2, while the complete list of original reactions is provided below.

**Fig. 2:**
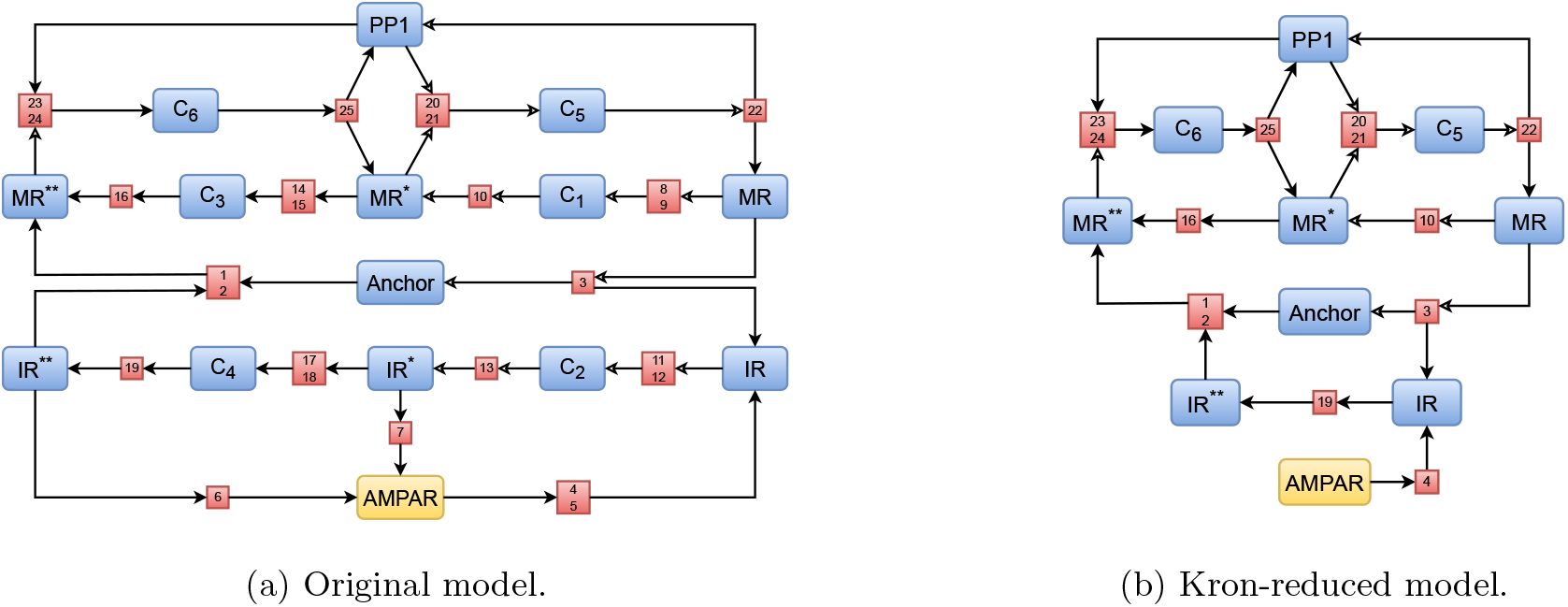
Schematic visualization of the original and Kron-reduced model of AMPAR trafficking cycle model with bistability. The left-hand panel shows the original model, as developed in [34]. The right-hand panel displays the Kron-reduced model, which is obtained by removing the species nodes {C_1_, C_2_, C_3_, C_4_, IR^*^} and reaction nodes {ℛ_*i*_ : *i* = 5, … 9, 11 …, 15, 17, 18} from the original graph. Nodes for non-buffered species are highlighted in blue, while the AMPAR node, as a buffered species, is in yellow, and reaction nodes appear in red.

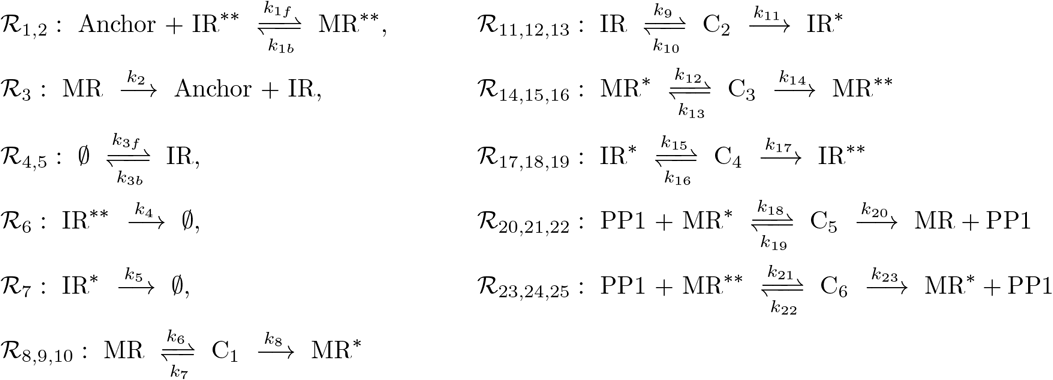

All reactions follow MAK, with reaction ℛ_4_ being influenced by the modifier AMPAR, while reactions ℛ_*i*_, *i* = 8, 11, 14, 17, are influenced by the modifier PKA. The model parameters and their respective values are listed in Table 1.

**Table 1:**
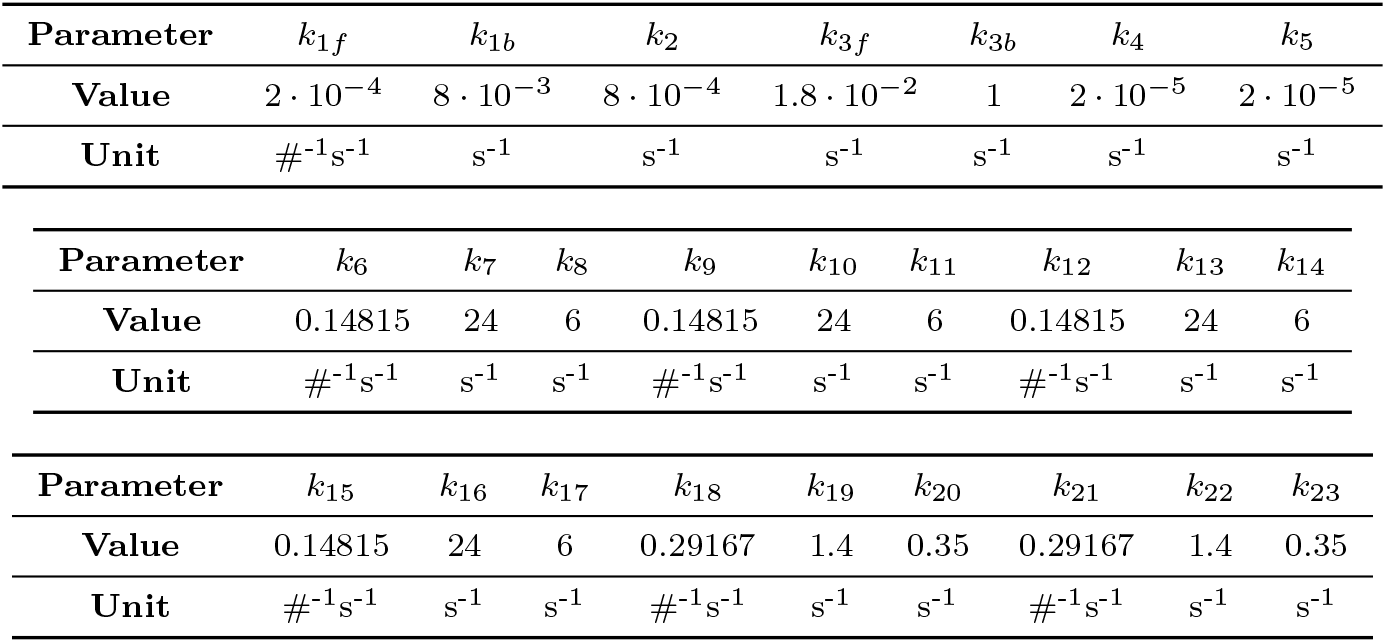
Parameter properties of the AMPAR trafficking cycle model.

Through our iterative process, the optimal combination of species and reactions for removal is identified, which includes the subset of species {C_1_, C_2_, C_3_, C_4_, IR^*^} and the subset of reactions {ℛ_*i*_ : *i* = 5, …, 9, 11, …, 15, 17, 18}. These subsets represent the largest group of nodes that can be removed while keeping the symmetrized error integral below 15%. The reduced model, therefore, consists of 9 species and 13 unidirectional reactions. The reduced CRN, along with its corresponding rates, is provided below.

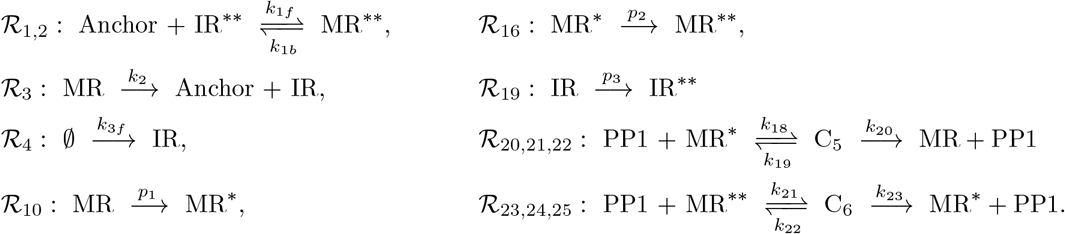

All reactions in the reduced model, similar to the original model, follow MAK, with reactions ℛ_10_, ℛ_16_, and ℛ_19_, influenced by the modifier PKA. The parameters *p*_1_, *p*_2_, and *p*_3_ are algebraic expressions of the original model parameters. Due to their complexity and length, explicitly presenting them in terms of the original parameters is impractical. However, their properties are summarized in Table 2.

**Table 2:**
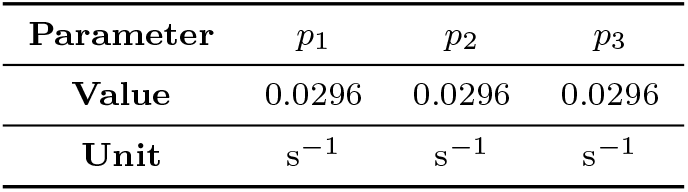
Parameters introduced in the reduced model that are absent in the original.

| Parameter | $p_1$ | $p_2$ | $p_3$ |
| --- | --- | --- | --- |
| Value | 0.0296 | 0.0296 | 0.0296 |
| Unit | $s^{-1}$ | $s^{-1}$ | $s^{-1}$ |

Fig. 3 presents a comparison of the evolution of molecular count of species between the reduced and original models for two distinct trajectories, both originating from the same stoichiometric compatibility class – that is, the set of all trajectories corresponding to the same conservation laws – but converging to different stable equilibrium points. Despite the removal of 35.71% of species and 48% of reactions, the discrepancy between the original and reduced models, as measured by the symmetrized error integral, remains relatively low. The error is 1.7334% for trajectories converging to one equilibrium state and 0.3377% for those reaching a different equilibrium state. Table 3 provides a quantitative comparison of the original AMPAR trafficking cycle model and the corresponding Kron-reduced model.

**Table 3:** Quantitative comparison of the original AMPAR trafficking cycle model and its reduced model.

|  | Species | Reactions | Deleted Species | Deleted Reactions | Error 1 | Error 2 |
| --- | --- | --- | --- | --- | --- | --- |
| Original model | 14 | 25 | — | — | — | — |
| Reduced model | 9 | 13 | 36% | 48% | 1.73% | 0.34% |

**Fig. 3:**
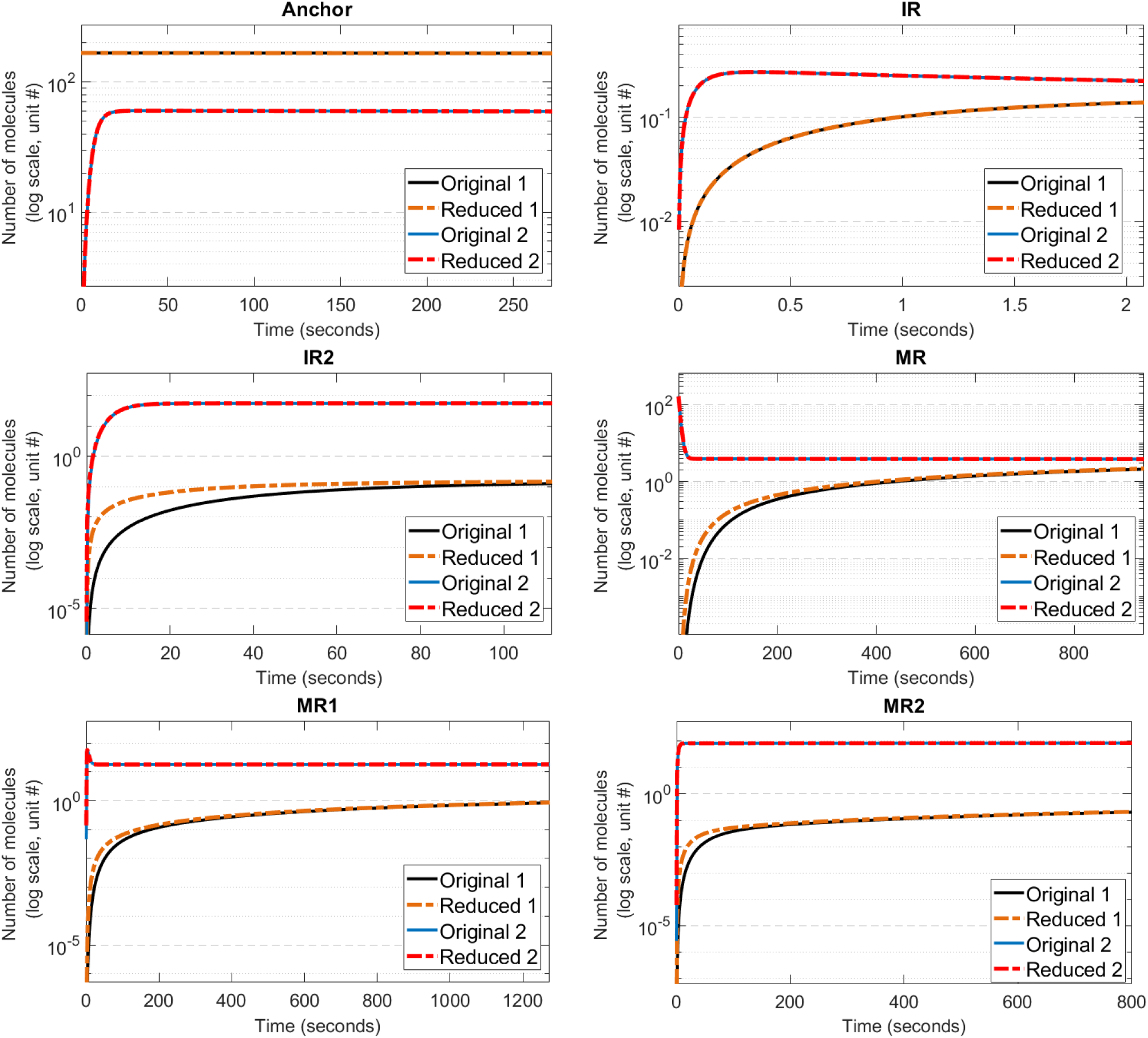
A comparison of species molecule numbers between the original and reduced AMPAR trafficking cycle models for trajectories converging to different stable equilibrium points.

## 6 Conclusion and discussion

High throughput methods for defining protein interaction and reaction maps [48–52] bring the possibility of constructing detailed and complex reaction models for important biological signaling networks such as in the immune system and neurobiology. While such models are necessary to simulate the effect of specific mutations and pharmacological interventions, they are computationally demanding and are harder to parameterize. Simpler but functionally equivalent models are easier to analyze and to parameterize. In this context it is desirable to fluently traverse between complex and functionally similar simple models. Thus, model reduction techniques transforming complex models into simpler models are useful tools in systems biology. Despite the importance of the problem, pipelines for automated model reduction have yet to be developed.

For these reasons, we proposed an automated model reduction method that is both flexible and easy to implement. While it shares some similarities with QSS and QE approximations [12, 14, 15] (as detailed in the supplementary material), there are important differences. QSS and QE methods determine which species and reactions to prune based on timescale and singular perturbation considerations. As discussed in [14, 15], these can be identified statically from the orders of magnitude of CRN parameters, without requiring simulations. However, the quality of the approximation is guaranteed only asymptotically, as timescale ratios (expressed as rational monomials of kinetic parameters) tend to zero.

In contrast, our reduction method simulates a large number of candidate reduced models and selects the best one. Its key advantage is its validity for finite timescale ratios. Another distinction lies in the pooling operations. QSS prunes fast species and replaces the reactions consuming and producing them with elementary modes, i.e., combinations of reactions that conserve the fast species and ensure direct exchanges between non-eliminated species [12]. The choice of these elementary modes is not necessarily unique. In contrast, our reduction method leads to a unique choice.

QE prunes fast reactions and pools species that are consumed or produced by them. The pools correspond to conservation laws of the subsystem of fast reactions [12, 15], and their choice may also be non-unique. In contrast, our reduction method prunes reactions whose effect on the total species concentration is negligible, without pooling species.

Note that our model reduction procedure involves simulations of both the original and reduced models in order to determine the optimal combination of nodes to be deleted and these simulations do pose a computational burden. However, this burden is insignificant compared with the burden of model analysis of the original model including stability/bifurcation analysis and parameter sensitivity analysis as described in the Introduction. Reducing the size of the model drastically reduces the computational efforts required to analyze the model. Moreover our model reduction procedure provides biological insights on which species or reactions of the original model play a dominant role in the dynamics and which ones have negligible influence since the ones with negligible influence get deleted through our procedure.

In the classical formulation of CRN dynamics based on the GCs [20], the Laplacian matrix is constant only under MAK. For GEKLs, however, the Laplacian may also depend on the system state through the reaction rate functions. In the SR-graph formulation considered here, the Laplacian depends explicitly on the state because the edge weights are defined via the reaction rates.

The original Kron reduction of GCs [20] is particularly effective for CRNs whose complex graphs contain connected components with more than two complexes (nodes). It reduces the number of complexes while retaining the remaining complexes and without adding new ones. This is biologically meaningful because the reduced network retains only complexes that already exist in the original biochemical mechanism, so every state in the reduced model corresponds to a real molecular configuration. Moreover, since no new complexes are introduced, the reduction avoids artificial biochemical states and preserves the interpretability of the CRN. However, when the GC consists of components with only two complexes, the original Kron reduction does not provide a meaningful reduction, as it may remove the entire reaction containing the complex we intend to remove. If such components share common species with other reactions, the extended Kron reduction [21, 22] can be applied. This two-step procedure first eliminates certain species using conservation laws, which can connect previously disconnected components of the GC into larger ones with multiple complexes; the original Kron reduction can then be meaningfully applied. A drawback of this approach is that it introduces additional parameters corresponding to the conservation constants, so the reduced model is valid only on trajectories consistent with these relations. As illustrated in the supplementary material with an example (see Section S1.2), the proposed Kron reduction of the SR-graph can, in certain cases, be dynamically equivalent to the extended Kron reduction of [22] as described above. For the considered example, both approaches lead to dynamically equivalent reduced models, but with different structures. On the other hand, compared with the Kron reduction methods based on the elimination of complexes [20–22], Kron reduction using SR-graphs is more flexible and leads to greater reduction corresponding to the same threshold. This has been illustrated using two examples of biomodels in Section S3 of the Supplementary Material. Indeed, the complex-based method cannot eliminate individual species, only entire complexes, which can be highly restrictive. We would like to remark that while the original complex-based Kron reduction approaches [20, 22] do preserve the kinetics of the original model, the SR-graph based Kron reduction approach introduced in this manuscript, may not do so. While the reduced model is still a valid chemical reaction network in terms of its network topology, it may not preserve the kinetics of the original.

The reduction of CRNs by making use of the graph Laplacian is facilitated by the well-established analogy between CRNs and electrical networks, based on shared structural, dynamical, and thermodynamical properties [42, 53]. Formally, the graph Laplacian is the negative generator of a diffusion process on the SR-graph [54]. In light of the above analogy, this process can be interpreted as the diffusion of information across the SR-graph. The associated diffusion flux respects the direction of causality: from reactions to the species they produce, and from species to the reactions whose rates they influence. Kron reduction enables the elimination of network elements (species and reactions) while preserving the essential pathways of information flow.

In summary, the proposed method offers several advantages, including a straightforward algorithmic implementation and uniquely defined SR-graph rewriting operations determined by the set of nodes to be pruned. Moreover, compared to Laplacian methods based on complexes, it provides greater flexibility by allowing the elimination of individual species or reactions rather than entire complexes. However, the approach has certain drawbacks, notably the need for simulations to validate the results and the risk of combinatorial explosion when identifying the nodes to prune, which may pose computational challenges.

The core procedure of our model reduction method, as discussed earlier, involves the removal of selected species and reaction nodes from the SR-graph associated with the CRN. This reduction is carried out by computing Schur complements of the Laplacian matrix corresponding to the SR-graph. We emphasize this aspect to highlight that the main reduction mechanism is independent of any specific trajectory of the system. However, the selection of species and reaction nodes to be removed is guided by the symmetrized error integral, which does depend on a base trajectory of the CRN.

Looking ahead, it is important to develop a trajectory-independent measure to quantify the discrepancy between the original and reduced models. This would enable a more robust and principled selection of species and reaction subsets for removal. One possible alternative to trajectory-based error measures could be the use of species and/or reaction settling times. Another alternative is to use orders of magnitude of parameters and tropical geometry methods to identify the nodes of the SR-graph to be pruned in the reduction method, along lines similar to those developed in [14, 15]. The proposed reduction method is formulated as a combinatorial optimization problem, in which admissible subsets of nodes are selected for elimination so as to minimize the symmetrized error integral. This problem is NP-hard. To manage its computational complexity, we currently limit the number of nodes that can be eliminated. As future work, we plan to improve our approach by implementing branch-and-bound strategies to efficiently explore the space of admissible node subsets. We also intend to use the reduction methods introduced in this paper to generate hierarchies of models with varying complexity, which are essential for hierarchical machine learning strategies [11].

## Ethics approval and consent to participate

Not applicable. This study did not involve human participants, animal subjects, or data requiring ethics approval.

## Consent for publication

Not applicable. This article does not contain any data, images, or information that requires consent for publication.

## Code availability

Python and MATLAB codes developed for this work are available at the following link: https://github.com/manvelgasparyan/model_reduction.

## Author contribution

SR formulated the initial concept. MG was responsible for formulating the problem, developing the solution, implementing the algorithm, and designing the automation tools. OR and SR contributed to the development of the solution and the main algorithms. USB and OR provided the models and biological insights, secured funding, and supervised the project. MG wrote the manuscript, with USB, OR and SR providing critical input, revisions, and review throughout the writing process.

## Acronyms

*AMPAR*: Activity-dependent Movement of a Glutamate Receptor.
*CDC*: Cell Division Cycle.
*CRN*: Chemical Reaction Network.
*DFG*: N-(1-deoxy-D-fructose-1-yl)-glycine.
*GC*: Graph of Complexes.
*GEKL*: General Enzyme Kinetics Law.
*HSP*: Hedgehog Signaling Pathway.
*MAK*: Mass-Action Kinetics.
*ODE*: Ordinary Differential Equation.
*PKA*: Protein Kinase A.
*QE*: Quasi-Equilibrium.
*QSS*: Quasi-Steady-State.
*SR-graph*: Species-Reaction Graph.

