## Supplementary Material for "Laplacian Dynamics and Kron Reduction in Species-Reaction Graphs of Chemical Reaction Networks"

Manvel Gasparyan, Upinder S. Bhalla, Ovidiu Radulescu, Shodhan Rao

#### S1 An illustrative simple example of the reduction procedure

We will illustrate the step-by-step process of reduction using a simple example of Chemical Reaction Network (CRN). Recall that our reduction approach assumes certain species are in a Quasi-Steady-State (QSS) condition, and that for specific reactions, there is no net change in any species resulting from these reactions. We will demonstrate the implications of these assumptions and outline the steps for reduction using the Schur complement of the Laplacian matrix.

##### S1.1 A simple mass-action chemical reaction network

Consider the following simple CRN of four species participating in one reversible and one irreversible reaction, governed by Mass-Action Kinetics (MAK):

$$\mathcal{S}_1 \underset{k_2}{\overset{k_1}{\rightleftharpoons}} \mathcal{S}_2, \qquad \mathcal{S}_2 + \mathcal{S}_3 \xrightarrow{k_3} \mathcal{S}_4, \tag{1}$$

with the rates given by:

$$v_1 = k_1 s_1, \qquad v_2 = k_2 s_2, \qquad v_3 = k_3 s_2 s_3,$$

where $k_i$ are positive constants. The Species-Reaction Graph (SR-graph) is visualized in the first panel of Fig. S1. The model represented by Eq. (12) of the main manuscript, pertaining to this example, is as follows:

$$\begin{aligned} \frac{ds_1}{dt} &= -k_1 s_1 + v_2, & 0 &= k_1 s_1 - v_1, \\ \frac{ds_2}{dt} &= v_1 - k_2 s_2 - k_3 s_2 s_3, & 0 &= k_2 s_2 - v_2, \\ \frac{ds_3}{dt} &= -k_3 s_2 s_3, & v_3 &= 2k_3 s_2 s_3 - v_3. \\ \frac{ds_4}{dt} &= v_3, \end{aligned} \tag{2}$$

Assuming that a selection criterion has led to the deletion of species $\mathcal{S}_2$ from the model. In line with our procedure, we assume that the species is in a QSS condition, i.e, $\frac{ds_2}{dt} = 0$. From this condition and the second equation of Eq. (2), we deduce:

$$s_2 = \frac{v_1}{k_2 + k_3 s_3}. \tag{3}$$

The directed graph obtained after deleting the node corresponding to the species $\mathcal{S}_2$ from the original SR-graph is visualized in the second panel of Fig. S1. As observed, this directed graph deviates from the structure of SR-graphs. However, it is noteworthy that this deviation can be rectified by deleting the nodes corresponding to reactions 1 and 2 from this directed graph. According to our procedure, this deletion is equivalent to setting the left-hand sides of the equations in the model (2), corresponding to reactions 1 and 2, to zero. Upon observation, we find that they are already zero since their corresponding stoichiometric

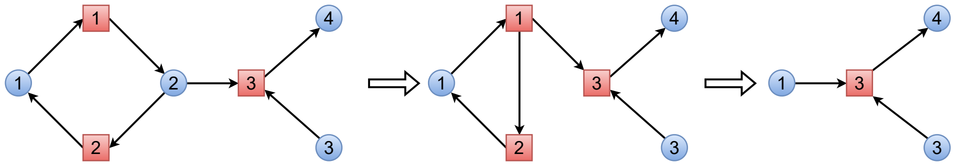


**Fig. S1**: Step-by-step reduction of the model of the CRN given in Eq. (2) is visualized in its SR-graph. First, species $\mathcal{S}_2$ is removed, creating direct links between reactions $\mathcal{R}_1$ and $\mathcal{R}_2$ with reaction $\mathcal{R}_3$. Next, reactions $\mathcal{R}_1$ and $\mathcal{R}_2$ are deleted, resulting in a direct connection from species $\mathcal{S}_1$ to $\mathcal{R}_3$.

imbalances are zero. Thus, it remains to derive $v_1$ and $v_2$ from these equations. From the aforementioned equations, we derive:

$$v_1 = k_1 s_1, \qquad v_2 = \frac{k_2 s_1}{k_2 + k_3 s_3},$$

and thus, we rewrite the model given in Eq. (2) as:

$$\begin{aligned} \frac{ds_1}{dt} &= -\frac{k_1 k_3 s_1 s_3}{k_2 + k_3 s_3}, & \frac{ds_4}{dt} &= v_3, \\ \frac{ds_3}{dt} &= -\frac{k_1 k_3 s_1 s_3}{k_2 + k_3 s_3}, & v_3 &= \frac{k_1 k_3 s_1 s_3}{k_2 + k_3 s_3}. \end{aligned} \tag{4}$$

The final obtained directed graph is visualized in the last panel of Fig. S1. As depicted, it now adheres to the structure of SR-graphs, thus representing a meaningful CRN. The final reduced model consists of a single irreversible reaction, shown below along with its reaction rate:

$$\mathcal{S}_1 + \mathcal{S}_3 \xrightarrow{v_3} \mathcal{S}_4, \qquad\qquad v_3 = \frac{k_1 k_3 s_1 s_3}{k_2 + k_3 s_3}. \tag{5}$$

It is evident that both the number of species and reactions have been reduced.

Let us present the reduction procedure using the Kron reduction formalism. It is straightforward to observe that, for this example, the vector of net stoichiometric imbalances is $\boldsymbol{d} = \begin{bmatrix} 0 & 0 & 1 \end{bmatrix}^\top$. Consequently, the model presented in Eq. (2) can be expressed in terms of the Laplacian matrix, similar to Eq. (12) of the main manuscript, as follows:

$$\begin{bmatrix} \frac{ds_1}{dt} \\ \frac{ds_2}{dt} \\ \frac{ds_3}{dt} \\ \frac{ds_4}{dt} \\ d_1 v_1 \\ d_2 v_2 \\ d_3 v_3 \end{bmatrix} = - \begin{bmatrix} k_1 & 0 & 0 & 0 & 0 & -1 & 0 \\ 0 & k_2 + k_3 s_3 & 0 & 0 & -1 & 0 & 0 \\ 0 & 0 & k_3 s_2 & 0 & 0 & 0 & 0 \\ 0 & 0 & 0 & 0 & 0 & 0 & -1 \\ -k_1 & 0 & 0 & 0 & 1 & 0 & 0 \\ 0 & -k_2 & 0 & 0 & 0 & 1 & 0 \\ 0 & -k_3 s_3 & -k_3 s_2 & 0 & 0 & 0 & 1 \end{bmatrix} \begin{bmatrix} s_1 \\ s_2 \\ s_3 \\ s_4 \\ v_1 \\ v_2 \\ v_3 \end{bmatrix}.$$

As in the previous case, let us eliminate the second species $\mathcal{S}_2$ along with the first and second reactions. Observe that $I = \{2, 5, 6\}$ denotes the set of indices corresponding to the second species, the first and second reactions, following the order of the SR-graph and Eq. (6). To proceed, we need to compute the Schur complement with respect to the set of indices $I$. For this, we consider the following block decomposition of the Laplacian matrix:

$$
\boldsymbol{L}_{11} = \begin{bmatrix} k_1 & 0 & 0 & 0 \\ 0 & k_3 s_2 & 0 & 0 \\ 0 & 0 & 0 & -1 \\ 0 & -k_3 s_2 & 0 & 1 \end{bmatrix}, \qquad \boldsymbol{L}_{12} = \begin{bmatrix} 0 & 0 & -1 \\ 0 & 0 & 0 \\ 0 & 0 & 0 \\ -k_3 s_3 & 0 & 0 \end{bmatrix},
$$

$$
\boldsymbol{L}_{21} = \begin{bmatrix} 0 & 0 & 0 & 0 \\ -k_1 & 0 & 0 & 0 \\ 0 & 0 & 0 & 0 \end{bmatrix}, \qquad \boldsymbol{L}_{22} = \begin{bmatrix} k_2 + k_3 s_3 & -1 & 0 \\ 0 & 1 & 0 \\ -k_2 & 0 & 1 \end{bmatrix}.
$$

In the next step, we compute the inverse of $\boldsymbol{L}_{22}$, which is given by:

$$
\boldsymbol{L}_{22}^{-1} = \begin{bmatrix} \dfrac{1}{k_2 + k_3 s_3} & \dfrac{1}{k_2 + k_3 s_3} & 0 \\ 0 & 1 & 0 \\ \dfrac{k_2}{k_2 + k_3 s_3} & \dfrac{k_2}{k_2 + k_3 s_3} & 1 \end{bmatrix}.
$$

Thus, the Schur complement of the Laplacian matrix $\boldsymbol{L}$ with respect to the set of indices $I$ is expressed as follows:

$$
\widehat{\boldsymbol{L}} = \boldsymbol{L}_{11} - \boldsymbol{L}_{12}\boldsymbol{L}_{22}^{-1}\boldsymbol{L}_{21} = \begin{bmatrix} \dfrac{k_1 k_3 s_3}{k_2 + k_3 s_3} & 0 & 0 & 0 \\ 0 & k_3 s_2 & 0 & 0 \\ 0 & 0 & 0 & -1 \\ -\dfrac{k_1 k_3 s_3}{k_2 + k_3 s_3} & -k_3 s_2 & 0 & 1 \end{bmatrix}.
$$

Using Eq. (3), we substitute the concentration of the eliminated species $s_2$ with the the corresponding function of the remaining variables and obtain

$$
\widehat{\boldsymbol{L}} = \begin{bmatrix} \dfrac{k_1 k_3 s_3}{k_2 + k_3 s_3} & 0 & 0 & 0 \\ 0 & \dfrac{k_1 k_3 s_1}{k_2 + k_3 s_3} & 0 & 0 \\ 0 & 0 & 0 & -1 \\ -\dfrac{k_1 k_3 s_3}{k_2 + k_3 s_3} & -\dfrac{k_1 k_3 s_1}{k_2 + k_3 s_3} & 0 & 1 \end{bmatrix}.
$$

It is important to note that, as stated in Theorem 1 of the main manuscript, the Schur complement $\widehat{\boldsymbol{L}}$ is again a Laplacian matrix. Consequently, the final Kron-reduced model can be expressed in terms of the Schur complement $\widehat{\boldsymbol{L}}$ as follows:

$$
\begin{bmatrix} \dfrac{ds_1}{dt} \\ \dfrac{ds_3}{dt} \\ \dfrac{ds_4}{dt} \\ d_3 v_3 \end{bmatrix} = - \begin{bmatrix} \dfrac{k_1 k_3 s_3}{k_2 + k_3 s_3} & 0 & 0 & 0 \\ 0 & \dfrac{k_1 k_3 s_1}{k_2 + k_3 s_3} & 0 & 0 \\ 0 & 0 & 0 & -1 \\ -\dfrac{k_1 k_3 s_3}{k_2 + k_3 s_3} & -\dfrac{k_1 k_3 s_1}{k_2 + k_3 s_3} & 0 & 1 \end{bmatrix} \begin{bmatrix} s_1 \\ s_3 \\ s_4 \\ v_3 \end{bmatrix},
$$

which leads to the identity given in Eq. (3) and the reduced model given in Eq. (5).

#### S1.2 Comparison with complex-based Kron reduction

In this section, we apply the Kron reduction method to the Graph of Complexes (GC) and compare it with our proposed reduction method, namely Kron reduction applied to the species–reaction graph, using the example given in (1). Our goal is to highlight the similarities and differences between the two reduction approaches.

First, we note that the original Kron reduction for GCs proposed in [1] cannot be applied directly, since the GC in this example is not connected, and removing any complex would also remove the corresponding reaction. For example, removing the complex $\mathcal{S}_2 + \mathcal{S}_3$ would eliminate the second reaction, leaving only the first reaction, which would substantially alter the system dynamics.

However, the extended Kron reduction method proposed in [2] can be applied. This approach first uses conservation laws to eliminate certain species, thereby connecting previously disconnected components into a single component, after which the original Kron reduction from [1] is performed.

In the present example, the method in [2] eliminates the species $\mathcal{S}_3$ using the conservation relation $s_3 = -s_4 + c$, where $c$ is a positive constant depending on the initial concentrations of $\mathcal{S}_3$ and $\mathcal{S}_4$. After this substitution, the network can be rewritten as $\mathcal{S}_1 \longleftrightarrow \mathcal{S}_2 \longrightarrow \mathcal{S}_4$ and Kron reduction is then applied to eliminate $\mathcal{S}_2$. This yields the reduced model

$$\mathcal{S}_1 \xrightarrow{\mathcal{S}_3} \mathcal{S}_4, \qquad s_3 = -s_4 + c, \qquad v_{\mathrm{r}} = \frac{k_1 k_3 s_1 s_3}{k_2 + k_3 s_3}, \qquad \frac{ds_1}{dt} = \frac{ds_4}{dt} = -v_{\mathrm{r}}. \tag{6}$$

We observe that Ordinary Differential Equations (ODEs) corresponding to the concentrations of the species $\mathcal{S}_1$ and $\mathcal{S}_4$ in this reduced model coincide with the ones obtained using our newly proposed method (see Section S1.1). The difference lies in the structure of the resulting reaction networks. In the reduction obtained via the method of [2], the species $\mathcal{S}_3$ appears indirectly through the conservation relation and acts as a modifier in the rate law. In contrast, in our new reduced model (5), $\mathcal{S}_3$ explicitly remains involved in the reaction stoichiometry, preserving its role in the original network.

The reduction method [2] introduces an additional parameter $c$, whose value is fixed for all trajectories. Consequently, the reduced model is valid only for trajectories that satisfy the corresponding conservation law for this fixed value of $c$. In contrast, when applying our new reduction method, the same conservation law appears in the reduced model, but the value of the conserved quantity is not fixed and instead varies depending on the trajectory. As a conclusion, for this example, the two reduction procedures mentioned above are dynamically equivalent but produce structurally different reduced reaction networks.

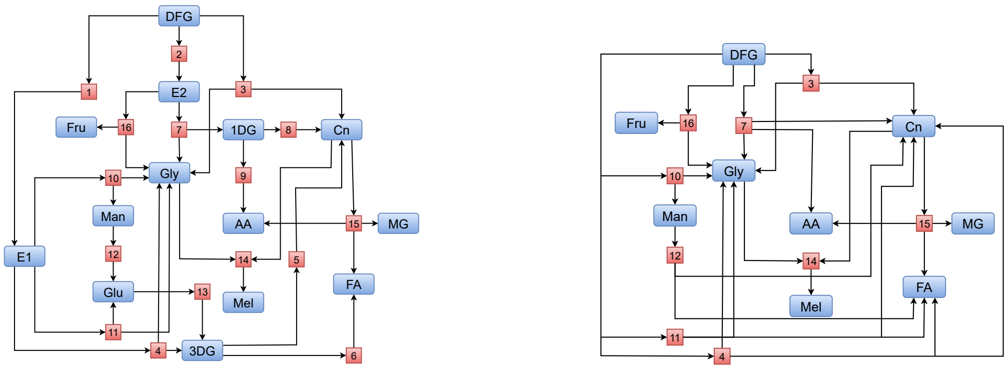


(a) Original SR-graph. (b) Kron-reduced SR-graph.

**Fig. S2**: Visualization of the SR-graphs for the original and Kron-reduced model of N-(1-deoxy-D-fructose-1-yl)-glycine thermal decomposition. The left-hand panel shows the original model, as developed in [3]. The right-hand panel displays the reduced SR-graph, which is obtained by removing the species nodes {E1, E2, 3DG, 1DG, Glu} and reaction nodes $\{\mathcal{R}_1, \mathcal{R}_2, \mathcal{R}_5, \mathcal{R}_6, \mathcal{R}_8, \mathcal{R}_9, \mathcal{R}_{13}\}$ from the original graph. Species nodes are highlighted in blue and reaction nodes appear in red.These figures depict a decrease in species and reactions from our reduction process, with a dynamic variation of 14.35% (symmetrized error integral).

| Parameter | $k_1$ | $k_2$ | $k_3$ | $k_4$ | $k_5$ | $k_6$ | $k_7$ | $k_8$ |
|---|---|---|---|---|---|---|---|---|
| **Value** | 0.0057 | 0.0156 | 0.0155 | 0.0794 | 0.0907 | 0.0274 | 0.2125 | 0.0000 |
| **Unit** | min$^{-1}$ | min$^{-1}$ | min$^{-1}$ | min$^{-1}$ | min$^{-1}$ | min$^{-1}$ | min$^{-1}$ | min$^{-1}$ |

| Parameter | $k_9$ | $k_{10}$ | $k_{11}$ | $k_{12}$ | $k_{13}$ | $k_{14}$ | $k_{15}$ | $k_{16}$ |
|---|---|---|---|---|---|---|---|---|
| **Value** | 1.9085 | 0.0707 | 0.1131 | 0.0008 | 0.0022 | 0.0034 | 0.0159 | 0.0134 |
| **Unit** | min$^{-1}$ | min$^{-1}$ | min$^{-1}$ | min$^{-1}$ | min$^{-1}$ | uM$^{-1}$ min$^{-1}$ | min$^{-1}$ | min$^{-1}$ |

**Table S1**: Parameter values of the original DFG thermal decomposition model.

#### S2 Reduction of a model of N-(1-deoxy-D-fructose-1-yl)-glycine

In addition to demonstrating our modeling and model reduction techniques on the Activity-dependent Movement of a Glutamate Receptor (AMPAR) trafficking cycle model [4] included in the main paper, we apply them to another real-life example of a CRN. We examine a mathematical model of N-(1-deoxy-D-fructose-1-yl)-glycine (DFG) thermal decomposition, as developed in [3] to investigate the specific reaction pathways, quantify the formation of key by-products like 3-Deoxyosone (3DG) and Acetic Acid (AA), and understand the effects of Potential of Hydrogen (pH) on these degradation processes. This model is sourced from the BioModels Database [5], accessible at https://www.ebi.ac.uk/biomodels/. Although the model is not overly complex, it captures the key features necessary to illustrate and deepen the understanding of our model reduction technique. The primary objective of this example is not to focus on reducing complexity, but rather to demonstrate the functionality and applicability of our method. The threshold value of the symmetrized error integral used to terminate the iterative reduction procedure was set to 0.15. This indicates that we are targeting a reduced model whose dynamics deviate by 15% from the original model's dynamics. In general, this threshold can be selected based on the desired level of similarity between the dynamics of the reduced model and those of the original model.

The process of DFG thermal decomposition is a complex process relevant to the Maillard reaction, which occurs when reducing sugars react with amino acids. A mathematical model has been developed to study this decomposition, focusing on how different pH levels influence the formation of key by-products such as 3DG, 1-Deoxyosone (1DG), and AA. The model provides insights into the specific reaction pathways, revealing that lower pH favors one degradation route while higher pH promotes another. This understanding not only enhances our knowledge of DFG breakdown but also aids in controlling chemical reactions in food and industrial applications, where these by-products can impact flavor, color, and quality.

The original system consists of 14 distinct chemical species interacting through 16 reactions governed by MAK, resulting in a SR-graph with 30 nodes. Fig. S2a provides a schematic representation of this graph, while the list of the reactions is provided below.

$$
\begin{aligned}
&\mathcal{R}_1: \mathrm{DFG} \xrightarrow{k_1} \mathrm{E1} && \mathcal{R}_9: \mathrm{1DG} \xrightarrow{k_9} \mathrm{AA}\\
&\mathcal{R}_2: \mathrm{DFG} \xrightarrow{k_2} \mathrm{E2} && \mathcal{R}_{10}: \mathrm{E1} \xrightarrow{k_{10}} \mathrm{Gly} + \mathrm{Man}\\
&\mathcal{R}_3: \mathrm{DFG} \xrightarrow{k_3} \mathrm{Gly} + \mathrm{Cn} && \mathcal{R}_{11}: \mathrm{E1} \xrightarrow{k_{11}} \mathrm{Gly} + \mathrm{Glu}\\
&\mathcal{R}_4: \mathrm{E1} \xrightarrow{k_4} \mathrm{Gly} + \mathrm{3DG} && \mathcal{R}_{12}: \mathrm{Man} \xrightarrow{k_{12}} \mathrm{Glu}\\
&\mathcal{R}_5: \mathrm{3DG} \xrightarrow{k_5} \mathrm{Cn} && \mathcal{R}_{13}: \mathrm{Glu} \xrightarrow{k_{13}} \mathrm{3DG}\\
&\mathcal{R}_6: \mathrm{3DG} \xrightarrow{k_6} \mathrm{FA} && \mathcal{R}_{14}: \mathrm{Gly} + \mathrm{Cn} \xrightarrow{k_{14}} \mathrm{Mel}\\
&\mathcal{R}_7: \mathrm{E2} \xrightarrow{k_7} \mathrm{Gly} + \mathrm{1DG} && \mathcal{R}_{15}: \mathrm{Cn} \xrightarrow{k_{15}} \mathrm{AA} + \mathrm{FA} + \mathrm{MG}\\
&\mathcal{R}_8: \mathrm{1DG} \xrightarrow{k_8} \mathrm{Cn} && \mathcal{R}_{16}: \mathrm{E2} \xrightarrow{k_{16}} \mathrm{Gly} + \mathrm{Fru}
\end{aligned}
$$

All the reactions are governed by MAK. The properties of the parameters involved in the model, including their values and units, are listed in Table S1.

We apply a systematic approach to reduce the model of DFG thermal decomposition by eliminating subsets of nodes containing exactly one species. First, we identify candidate subsets of species and reaction nodes with a species portion $N = 1$. These candidate subsets are ranked according to the values of the symmetrized error integral defined in Eq. (8) of the main manuscript., and the subset with the smallest value is removed. This process is then applied iteratively to the reduced model, repeating until the symmetrized error integral reaches a predefined threshold, which in this case is set to 0.15. This procedure identifies sets

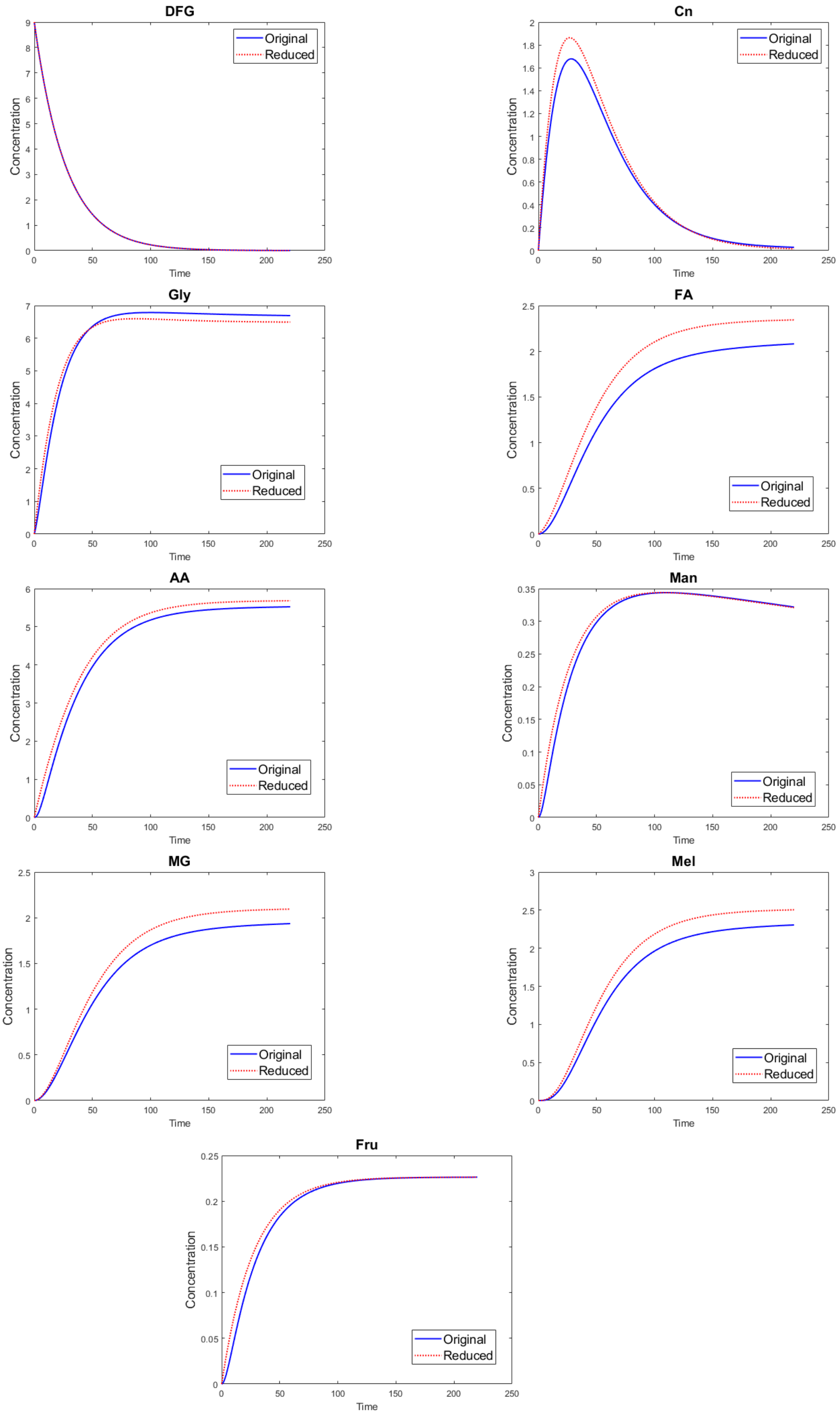


**Fig. S3**: Comparison of species concentrations in the N-(1-deoxy-D-fructose-1-yl)-glycine thermal decomposition model between the original model and reduced model.

| Parameter | $p_4$ | $p_7$ | $p_{10}$ | $p_{11}$ | $p_{16}$ |
|---|---|---|---|---|---|
| **Expression** | $\frac{k_1k_4}{k_4+k_{10}+k_{11}}$ | $\frac{k_2k_7}{k_2+k_7}$ | $\frac{k_1k_{10}}{k_4+k_{10}+k_{11}}$ | $\frac{k_1k_{11}}{k_4+k_{10}+k_{11}}$ | $\frac{k_2k_{16}}{k_7+k_{16}}$ |
| **Value** | 0.0017 | 0.0147 | 0.0015 | 0.0024 | $9.25\times10^{-4}$ |
| **Unit** | min$^{-1}$ | min$^{-1}$ | min$^{-1}$ | min$^{-1}$ | min$^{-1}$ |

**Table S2**: Formulas used to determine the parameters of the reduced DFG thermal decomposition model.

{E1, E2, 3DG, 1DG, Glu}, and $\{\mathcal{R}_i : i = 1, 2, 5, 6, 8, 9, 13\}$ as the optimal combination of species and reactions for removal, representing the largest subset of nodes with a symmetrized error integral of no more than 15%. Deleting these nodes results in the reduced network, with the corresponding SR-graph shown in Fig. S2b. Using Eq. (13) in Remark 2 of the main manuscript., we determine the reaction rates for the reactions in the Kron-reduced model. The complete list of reactions and their corresponding reaction rates in the final Kron-reduced model is provided below.

$$\mathcal{R}_3 : \text{DFG} \xrightarrow{k_3} \text{Gly} + \text{Cn} \qquad \mathcal{R}_{11} : \text{DFG} \xrightarrow{p_{11}} \text{Gly} + \text{FA}$$

$$\mathcal{R}_4 : \text{DFG} \xrightarrow{p_4} \text{Gly} + \text{Cn} + \text{FA} \qquad \mathcal{R}_{12} : \text{Man} \xrightarrow{k_{12}} \text{Cn} + \text{FA}$$

$$\mathcal{R}_7 : \text{DFG} \xrightarrow{p_7} \text{Gly} + \text{Cn} + \text{AA} \qquad \mathcal{R}_{14} : \text{Gly} + \text{Cn} \xrightarrow{k_{14}} \text{Mel}$$

$$\mathcal{R}_{10} : \text{DFG} \xrightarrow{p_{10}} \text{Gly} + \text{Man} \qquad \mathcal{R}_{15} : \text{Cn} \xrightarrow{k_{15}} \text{AA} + \text{FA} + \text{MG}$$

$$\mathcal{R}_{16} : \text{DFG} \xrightarrow{p_{16}} \text{Gly} + \text{Fru}$$

The comparison of key species concentrations between the original and reduced models is illustrated in Fig. S3. Notably, despite the elimination of 35.71% of species and 43.75% of reactions, the resulting error integral is only 14.35%. Table S3 provides a quantitative comparison of the original model of DFG and the corresponding Kron-reduced model.

| | Species | Reactions | Deleted Species | Deleted Reactions | Error Integral |
|---|---|---|---|---|---|
| **Original model** | 14 | 16 | — | — | — |
| **Reduced model** | 9 | 9 | 35.71 % | 43.75 % | 14.25 % |

**Table S3**: Quantitative comparison of the original model and the reduced model for the DFG thermal decomposition model.

### S3 Additional case studies

In this section, we further illustrate the applicability of the proposed model reduction method using two additional real-world examples of CRNs. The models considered here are taken from the BioModels Database [5]. Unlike the previous examples, which were analyzed in detail to demonstrate the methodology step by step, the purpose here is to compare the application of the method of Kron reduction on SR-graphs introduced in this manuscript with the application of Kron reduction on the GC as described in [1, 2].

#### S3.1 Cell division cycle: cdc2 and cyclin interactions

We consider a model of the Cell Division Cycle (CDC) describing the interaction between cdc2 and cyclin, as developed in [6]. The model consists of six species and eight reactions. The original SR-graph and GC are shown in Fig. S4a and Fig. S4c, respectively. Although the model is relatively small in terms of the number of species, reactions, and the corresponding system of ODEs – so that model reduction would not typically be required – it nevertheless provides a convenient example for illustrating how both Kron reduction approaches operate and how they compare. The aim here is not to achieve further simplification, but rather to demonstrate how these two reductions are performed on a real-life example.

We first apply the proposed reduction approach based on Kron reduction of the SR-graph associated with the model. This procedure eliminates the species $\mathcal{S}_1$ and $\mathcal{S}_4$, together with the reactions $\mathcal{R}_2, \mathcal{R}_3, \mathcal{R}_5, \mathcal{R}_8$, resulting in a reduced CRN whose corresponding SR-graph is shown in Fig. S4b. Next, we perform Kron reduction on the GC of the model. Following the procedure described in [1], this reduction removes the complex $\mathcal{S}_4$ as well as the empty complex, which represents the inflow and outflow of the network.

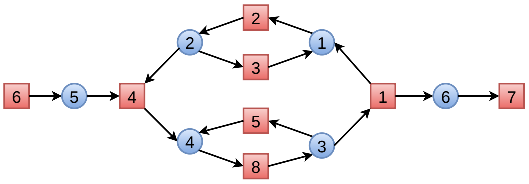

(a) Original SR-graph.

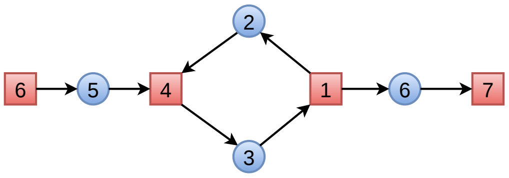

(b) Kron-reduced SR-graph.

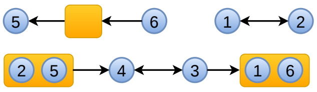

(c) Original GC.

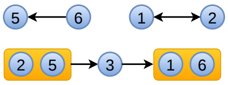

(d) Kron-reduced GC.

**Fig. S4**: Visualization of the SR-graphs and GCs for the original and Kron-reduced models of the CDC. Species are highlighted in blue, reactions in red, and complexes in orange.

**Table S4**: Comparison of the original model with the two reduced models obtained via Kron reduction based on the SR-graph and the GC for the model of CDC.

| Model | Species | Reactions | Deleted Species | Deleted Reactions | Error Integral |
|---|---|---|---|---|---|
| **Original Model** | 6 | 8 | — | — | — |
| **Reduced Model** (SR-graph) | 4 | 4 | 33 % | 50 % | 12.23% |
| **Reduced Model** (GC) | 5 | 5 | 16 % | 37 % | 11.55% |

Table S4 summarizes the structural reductions and the corresponding dynamical discrepancies for both approaches. As can be seen, the newly proposed Kron reduction applied to the SR-graph removes two species, reducing the number of species from six to four, which corresponds to a 33.33% reduction. It also decreases the number of reactions from eight to four, corresponding to a 50% reduction. In contrast, the original Kron reduction based on the GC removes only one species, decreasing the number of species from six to five and reducing the number of reactions from eight to five. This corresponds to a 16.67% reduction in species and a 37.5% reduction in reactions. Despite these structural changes, the dynamical discrepancy between the original and the reduced models remains relatively small: 12.13% for the reduction based on the SR-graph and 11.55% for the reduction based on the GC. Therefore, the proposed SR-graph-based reduction achieves a substantially greater structural simplification while maintaining a comparable level of dynamical accuracy, making it a more effective approach for reducing the complexity of the model.

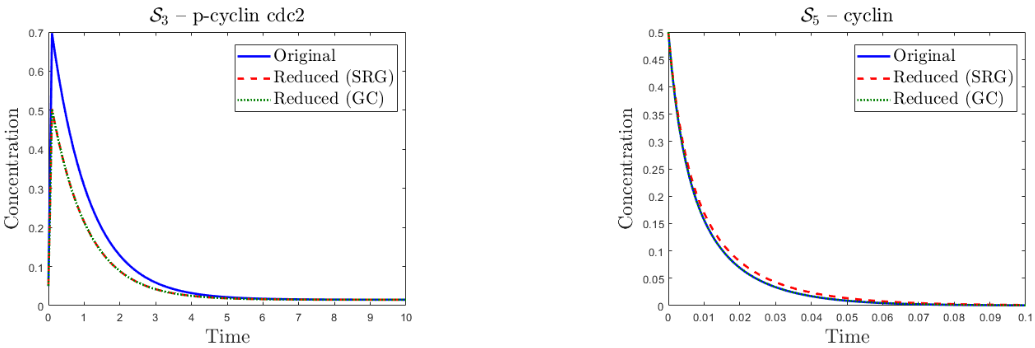


**Fig. S5**: Comparison of species concentrations in the original CDC model and the corresponding Kron-reduced models.

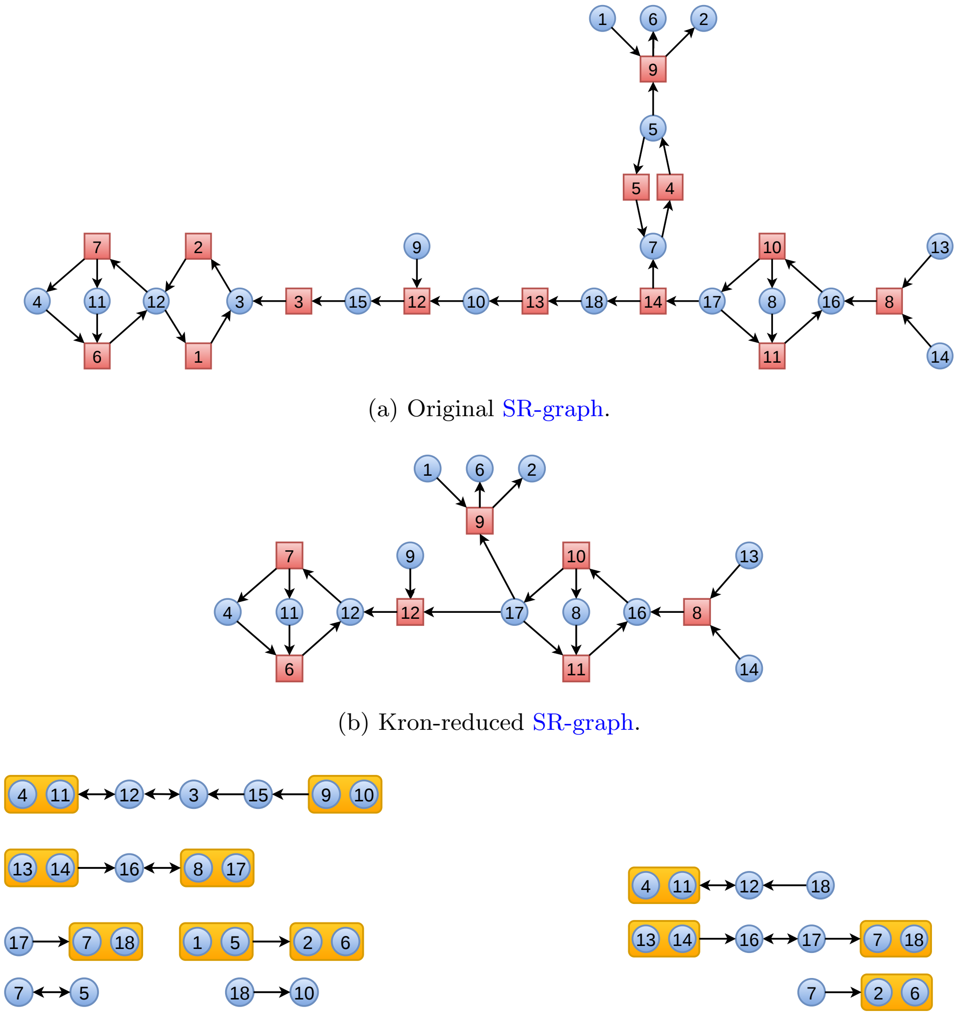

(a) Original SR-graph.

(b) Kron-reduced SR-graph.

(c) Original GC.

(d) Kron-reduced GC.

**Fig. S6**: Visualization of the SR-graphs and GCs for the original and Kron-reduced models of the HSP. Species are highlighted in blue, reactions in red, and complexes in orange.

#### S3.2 Hedgehog signalling pathway

We applied our new model reduction technique to a different type of CRN, namely a model of the Hedgehog Signaling Pathway (HSP). The detailed mathematical formulation of this network is available in [7]. The original CRN consists of 14 irreversible reactions involving 18 species. The original SR-graph and the original GC are shown in Fig. S6a and Fig. S6c, respectively.

We first apply our newly proposed model reduction method, which performs Kron reduction on the SR-graph of this model. After applying the reduction, species $S_3, S_5, S_7, S_{10}, S_{15}, S_{18}$ and reactions $\mathcal{R}_1, \mathcal{R}_2, \mathcal{R}_3, \mathcal{R}_4, \mathcal{R}_5, \mathcal{R}_{13}, \mathcal{R}_{14}$ are removed, yielding a reduced CRN whose corresponding SR-graph is shown in Fig. S6b. For this model, the Kron reduction on the GC with conservation laws [2] gives better reduction than the original Kron reduction [1] on GC due to small connected components in the GC. Therefore, the comparison in this case is with the method of [2]. The original GC is shown in Fig. S6c. The first step in the method of [2] leads to the elimination of the species $\mathcal{S}_1, \mathcal{S}_8, \mathcal{S}_9$ from the model using conservation laws. The second step of this method is the Kron reduction of the resulting GC, which leads to the removal of the complexes $\mathcal{S}_3, \mathcal{S}_5, \mathcal{S}_{10}, \mathcal{S}_{15}$ from the model.

The sizes of the original and reduced models obtained by the two reduction approaches are summarized in Table S5. Recall that the original model consists of 18 species and 14 reactions.

Both approaches achieve a substantial simplification of the original network while preserving its main dynamical features. The reduction based on the SR-graph decreases the model from 18 to 12 species (a reduction of about 33%) and from 14 to 7 reactions (a 50% reduction), while maintaining a small approximation error of 5.27%. In contrast, the reduction based on the GC yields a model with 13 species (about 28% fewer species) and 8 reactions (about 43% fewer reactions) yielding an approximation error of about

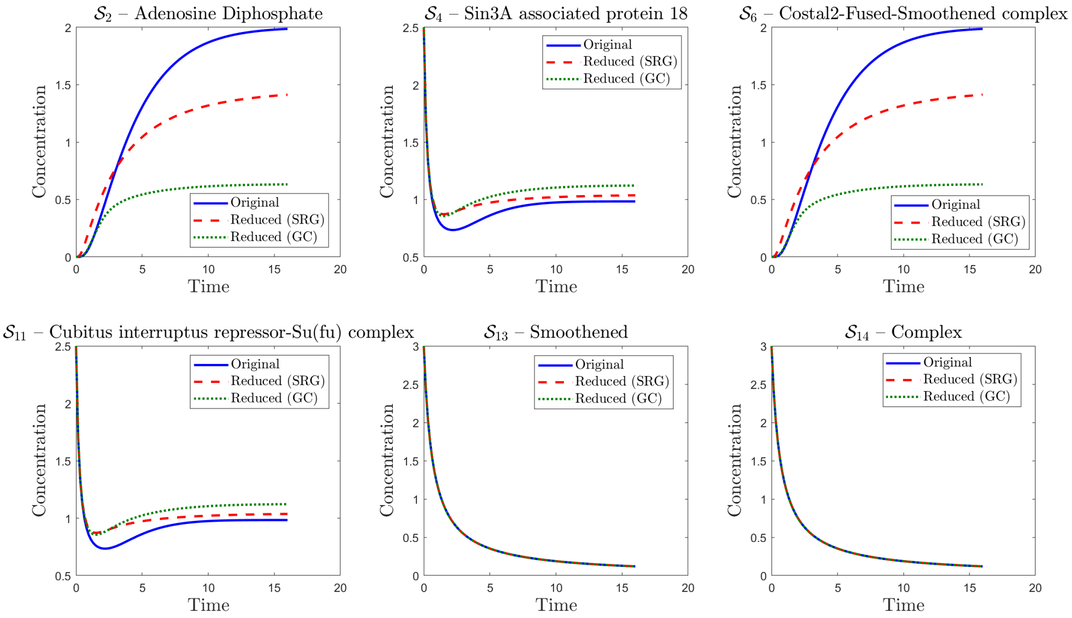


**Fig. S7**: Comparison of species concentrations in the original HSP model and the corresponding Kron-reduced models.

6.59%. A comparison between the concentrations of several remaining species in the original model and the reduced models is shown in Fig. S7.

The reduction based on the SR-graph produces a slightly more compact model, removing more species and reactions while maintaining a small approximation error. This makes it advantageous when the primary objective is to reduce the dimensionality of the system at the level of species dynamics. In contrast, the reduction based on the GC operates at the level of complexes and is more closely aligned with the structural organization of the CRN. As a result, it may better preserve certain structural properties of the CRN, which can be beneficial for theoretical analysis. On the other hand, the Kron reduction on the GC introduces additional parameters in the reduced model that depend on the initial conditions of the species, i.e., on the base trajectories. Consequently, the resulting reduced model is trajectory-dependent and may not be valid for arbitrary initial conditions.

**Table S5**: Comparison of the original model with the two reduced models obtained via Kron reduction based on the SR-graph and the GC for the model of HSP.

| Model | Species | Reactions | Deleted Species | Deleted Reactions | Error Integral |
|---|---|---|---|---|---|
| **Original Model** | 18 | 14 | — | — | — |
| **Reduced Model** (SR-graph) | 12 | 7 | 33 % | 50 % | 5.27% |
| **Reduced Model** (GC) | 13 | 8 | 28 % | 43 % | 6.59% |

### S4 Relation with quasi-steady state and quasi-equilibrium approximation based reductions

QSS and Quasi-Equilibrium (QE) approximations are two widely used methods for reducing CRNs. While some preliminary theoretical results are available in [8], the algorithmic implementation of these methods has not yet been fully developed. we revisit these reduction algorithms and compare them with our proposed approach presented in the main manuscript—namely, the Kron reduction of CRNs based on the SR-graph and its associated Laplacian dynamics.

The first, more challenging part of QSS and QE reduction methods is the identification of a fast subsystem (comprising fast species and reactions). This is achieved using the geometric theory of singular perturbations and tropical geometry-based scalings [9, 10]. Although these reduction methods have been described algorithmically, the cited works apply the geometric theory of singular perturbations algebraically to the system of ODEs governing the CRN dynamics. The method involves eliminating fast species variables and introducing new pseudo-species variables, which are linear or nonlinear functions of species concentrations, approximately conserved by the fast reactions (approximate conservation laws, see [10]). These eliminations and additions are applied algebraically to the set of ODEs. The SR-graph structure is not easily recognizable in the final reduced set of ODEs.

However, in [8] a graph theoretical approach is provided for QSS and QE. In this approach, the QSS and QE reductions are formulated as graph rewriting operations, transforming a SR-graph into a simplified SR-graph. BothQSS and QE operations result from two types of operations: species or reaction pruning in which species or reactions are eliminated and species or reaction pooling in which species and reactions are combined to form new species and reactions. This graphical approach is closely related to the Kron reduction of the SR-graph, which involves pruning certain species and reactions, and redefining others.

In the following section, we mention fast species and reactions without specifying how they were identified. The QSS and QE reductions are then applied formally. For the detection of fast subsystems, we refer to [9, 10].

#### S4.1 Comparison of QSS and Kron reduction

The QSS reduction method consists in pruning a number of fast species and combining reactions into linear combinations (reaction pools) that leave the fast species unchanged. The number of reaction pools is smaller than the number of pooled reactions, therefore QSS reduces not only the number of species but also the number of reactions.

The QSS reduction is formalized as follows (see [8]). Let $(\mathcal{V}_s, \mathcal{V}_r, \mathcal{E})$ be the SR-graph, and $I \subset \mathcal{V}_s$ denote the subset of fast species to be pruned. The species in the complementary subset $T = \mathcal{V}_s \setminus I$ are called terminal and are retained in the reduced chemical reaction network model.

To $I$, one associates a reaction subset $R_I \subset \mathcal{V}_r$ of all the reactions that either produce, or consume species from $I$. Let $\boldsymbol{S}_I \in \mathbb{Z}^{n_I} \times \mathbb{Z}^{r_I}$ ($n_I, r_I$ are the number of species in $I$ and the number of reactions in $R_I$, respectively) be the stoichiometric matrix of reactions from $R_I$, projected on the directions of species $I$, i.e. a matrix whose columns are the vectors $[\beta_{ij} - \alpha_{ij}]_{i \in I}$, $j \in R_I$. We also define $\boldsymbol{S}_T \in \mathbb{Z}^{n_T} \times \mathbb{Z}^{r_I}$ ($n_T$ is the number of species in $T$) as the terminal stoichiometric matrix whose columns correspond to the reactions $R_I$, representing the action of these reactions on terminal species $[\beta_{ij} - \alpha_{ij}]_{i \in T}$, $j \in R_I$.

With these definitions, the stoichiometric matrix of the CRN has the following block decomposition:

$$\boldsymbol{S} = \begin{bmatrix} \boldsymbol{S}_I & \mathbb{O} \\ \boldsymbol{S}_T & \boldsymbol{S}_E \end{bmatrix},$$

where $\boldsymbol{S}_E$ contains the stoichiometries of external reactions $V_r \setminus R_I$.

The reactions $R_I$ are then pooled into *elementary modes* with non-zero terminal stoichiometry. An elementary mode [11] is a linear combination of stoichiometric vectors of reactions $R_I$, with coefficients $\boldsymbol{\gamma} = [\gamma_1, \gamma_2, \ldots, \gamma_{r_I}]^T$ satisfying the following properties:

*i)* species in $I$ are stationary under the action of such combinations,

$$\boldsymbol{S}_I \boldsymbol{\gamma} = 0. \tag{7}$$

*ii)* irreversible reactions have positive coefficients in the combination, $\gamma_j \geq 0$ if reaction $j$ is irreversible.
*iii)* the combinations are minimal, in the sense that if $\boldsymbol{\gamma} = \boldsymbol{\gamma}' + \boldsymbol{\gamma}''$ where $\boldsymbol{\gamma}'$ and $\boldsymbol{\gamma}''$ satisfy i),ii) and have more zero components than $\boldsymbol{\gamma}$, then $\boldsymbol{\gamma}' = 0$ or $\boldsymbol{\gamma}'' = 0$.

We add to these conditions the following condition

*iv)* the combinations have non-zero terminal stoichiometry

$$\boldsymbol{S}_T \boldsymbol{\gamma} \neq 0. \tag{8}$$

According to conditions i)-iii) the elementary modes are minimal circuits, or matroid bases of reactions in $R_I$. In the framework of this paper all reactions are considered irreversible or split into a forward and reverse irreversible reaction when they are reversible. In this case, elementary modes are equivalent to extreme vectors (convex basis of the cone defined by the conditions i) and ii)) [12, 13]. Let us remind that a vector is extreme if it satisfies the conditions i) and ii) and are minimal generators of the cone described by i) and ii) (minimality meaning that they cannot be expressed by combinations with positive coefficients of two or more vectors in the cone). Elementary modes and extreme vectors were well studied in the stoichiometry theory of CRNs and several computational tools implement algorithms for computing them [11, 14–16].

To compute the reaction rates of the reaction pools, we need to ensure that the contributions of reactions from $R_I$ to the time derivatives of temporal species are equal before and after pooling, meaning that

$$\boldsymbol{S_T v}_I(\boldsymbol{s}) = \sum_{i=1}^{m_I} \hat{v}_i \boldsymbol{S}_T \boldsymbol{\gamma}_i, \tag{9}$$

where $\boldsymbol{\gamma}_i$ are the extreme vectors with non-zero terminal stoichiometries, $m_I$ is the number of these vectors, $\hat{v}_i$ are the reaction rates to be computed, $\boldsymbol{v}$ is the vector of reaction rates of the full model [8].

We also need to express the concentrations of pruned species as functions of concentrations of terminal species by solving the QSS equations:

$$\boldsymbol{S_I v}_I(\boldsymbol{s}) = 0, \tag{10}$$

where $\boldsymbol{v}_I$ are the rates of the reactions $R_I$.

The species elimination is possible under the following necessary condition:

*v)* $\text{rank}(\boldsymbol{S_I}) = n_I$.

Furthermore, Eq. (9) has a unique solution under the following condition:

*vi)* The vectors $\boldsymbol{S}_T\boldsymbol{\gamma}_i$ are linearly independent (i.e., the extremal vectors have independent terminal stoichiometries).

In Kron reduction, degeneracy of the Schur complement can also occur when the Laplacian diagonal block $\mathbf{L}_{22}$ is singular; however, this condition is generally different from the condition (vi). The quasi-steady-state equations (10), and the limitations that follow from them, are exactly the same in both QSS and Kron reduction (see Remark 1 in the main text).

It is also possible that eq. (9) has no solution. The most common case arises when there are no extreme vectors $\boldsymbol{\gamma}_i$ satisfying condition (iv). In general, (9) has solutions if and only if:

*vii)* $\text{rank}([\boldsymbol{S_T v}_I(\boldsymbol{s}), \boldsymbol{S}_T\boldsymbol{\gamma}_1, \ldots, \boldsymbol{S}_T\boldsymbol{\gamma}_{m_I}]) = \text{rank}([\boldsymbol{S}_T\boldsymbol{\gamma}_1, \ldots, \boldsymbol{S}_T\boldsymbol{\gamma}_{m_I}]) = m_I$.

In summary, the QSS reduction algorithm is as follows

---

**Algorithm S1** QSSReduction

---

**Require:** A CRN given by $\boldsymbol{S}$ matrix and reaction rate vector $\boldsymbol{v}(\boldsymbol{s})$. A subset of species $I$ to prune.

**Ensure:** A reduced CRN with less reactions and less species, given by a smaller $\hat{\boldsymbol{S}}$ matrix and reaction rate vector $\hat{\boldsymbol{v}}(\hat{\boldsymbol{s}})$.

1: Compute the sets $I$, $R_I$, $R_T$, and the matrices $\boldsymbol{S}_I$, $\boldsymbol{S}_T$, $\boldsymbol{S}_E$.

2: Eliminate the species I.

3: Compute $\boldsymbol{\gamma}_j$, $1 \leq j \leq m_I \leq r_I$ satisfying i)-iv).

4: Solve (10) for $\boldsymbol{s}_I$.

5: Solve (9) for $\hat{v}_j$.

6: Use solution of (10) obtained at **3.** to express $\hat{v}_j$, and $\boldsymbol{v}_T$ as functions of $\boldsymbol{s}_I$ and $\boldsymbol{s}_T$.

7: Define the new reactions: $m_I$ reaction pools with stoichiometries $\boldsymbol{S_T}\boldsymbol{\gamma}_j$, and rates $\hat{v}_j$; the reactions $R_T$ with stoichiometries $\boldsymbol{S}_E$ and rates $\boldsymbol{v}_T$.

8: Define $\hat{\boldsymbol{S}} = [\boldsymbol{S_T}\boldsymbol{\gamma}_1, \ldots, \boldsymbol{S_T}\boldsymbol{\gamma}_{m_I}, \boldsymbol{S}_E]$, $\hat{\boldsymbol{v}} = \begin{bmatrix} \hat{v}_1 \\ \vdots \\ \hat{v}_{m_I} \\ \hat{\boldsymbol{v}}_T \end{bmatrix}$, $\hat{\boldsymbol{s}} = \boldsymbol{s}_T$.

---

To illustrate these concepts, let us revisit the example presented in the Section S1. In this example there is only one species to eliminate $I = \{2\}$ and three terminal species to retain $T = \{1, 3, 4\}$. All the three reactions of the model consume or produce the eliminated species, namely $R_I = \{1, 2, 3\}$. These define

$$\boldsymbol{S}_I = \begin{bmatrix} 1 & -1 & -1 \end{bmatrix}, \; \boldsymbol{S}_T = \begin{bmatrix} -1 & 1 & 0 \\ 0 & 0 & -1 \\ 0 & 0 & 1 \end{bmatrix},$$

and $\boldsymbol{S}_E$ is an empty matrix.

$\boldsymbol{S}_I$ has two extreme vectors $\begin{bmatrix} 1 & 0 & 1 \end{bmatrix}^\top$, $\begin{bmatrix} 1 & 1 & 0 \end{bmatrix}^\top$ but the second one has zero terminal stoichiometry and must be excluded. The reduced model consists in eliminating the species 2 and replacing the three reactions involving this species by the combination $[101]^T$ that has terminal stoichiometry $[-1-11]^T$ (it consumes species $1, 3$ and produces the species 4). The reduced model has only one reaction corresponding to the extreme vector $[101]^T$ and the stoichiometry matrix of the reduced model reads

$$\boldsymbol{S}_{red} = \boldsymbol{S}_T \begin{bmatrix} 1 \\ 0 \\ 1 \end{bmatrix} = \begin{bmatrix} -1 \\ -1 \\ 1 \end{bmatrix}.$$

The vector of reaction rates is $\boldsymbol{v} = [k_1 s_1, k_2 s_2, k_3 s_2 s_3]^T$. From (10) we get $s_2 = k_1 s_1/(k_2 + k_3 s_3)$. Eq.(9) leads to

$$\begin{bmatrix} -k_1 s_1 + k_2 s_2 \\ -k_3 s_2 s_3 \\ k_3 s_2 s_3 \end{bmatrix} = \frac{k_1 k_3 s_1 s_3}{k_2 + k_3 s_3} \begin{bmatrix} -1 \\ -1 \\ 1 \end{bmatrix} = \hat{v}_1 \begin{bmatrix} -1 \\ -1 \\ 1 \end{bmatrix},$$

and therefore the reaction rate of the only reaction of the reduced model is

$$\hat{v}_1 = \frac{k_1 k_3 s_1 s_3}{k_2 + k_3 s_3}.$$

The reduced model is composed of a single reaction $S_1 + S_3 \to S_4$, with $\hat{g}_1 = \frac{k_1 k_3}{k_2 + k_3 s_3}$. In this case the QSS reduction is completely equivalent to the Kron reduction (see (5) and Figure S1). Let us also present situations where Kron reduction and QSS reductions differ. To this end, we revisit the Example 1 from the main text.

The stoichiometric matrix of the full model is:

$$S = \begin{bmatrix} -1 & 1 & 2 \\ -2 & 2 & 0 \\ 3 & -3 & 0 \\ 1 & -1 & -4 \\ 0 & 0 & 5 \end{bmatrix}$$

Let us consider first that we want to eliminate species 1 and 2. In this case, the set $R_I$ contains all the reactions and

$$\boldsymbol{S}_I = \begin{bmatrix} -1 & 1 & 2 \\ -2 & 2 & 0 \end{bmatrix}, \ \boldsymbol{S}_T = \begin{bmatrix} 3 & -3 & 0 \\ 1 & -1 & -4 \\ 0 & 0 & 5 \end{bmatrix},$$

and $\boldsymbol{S}_E$ is an empty matrix. Eq. (7) defining extreme vectors (condition i)) has one dimensional solutions $\gamma = \alpha(1, 1, 0)^T$ such that $\boldsymbol{S}_T \gamma = 0$. Conditions vi) and vii) are not satisfied, therefore the QSS reduction is not possible.

Consider now the case when we prune only the species 1, $I = \{1\}$. In this case,

$$\boldsymbol{S}_I = \begin{bmatrix} -1 & 1 & 2 \end{bmatrix}, \ \boldsymbol{S}_T = \begin{bmatrix} -2 & 2 & 0 \\ 3 & -3 & 0 \\ 1 & -1 & -4 \\ 0 & 0 & 5 \end{bmatrix},$$

and $\boldsymbol{S}_E$ is empty.

The equation (10) has now two dimensional solutions $\gamma = \alpha(1, 1, 0)^T + \beta(2, 0, 1)^T$. There are two extreme vectors but only one, $\gamma = (2, 0, 1)^T$, satisfies (8).

To compute the reaction rate of the reduced reaction we use (9) in combination with (10). These read:

$$\begin{aligned} -4\hat{v} &= -2v_1 - 2v_2, \\ 6\hat{v} &= v_1 - v_2 - 4v_3, \\ -2\hat{v} &= 3v_1 - v_2, \\ 5\hat{v} &= 5v_3, \\ v_1 &= v_2 + 2v_3, \end{aligned}$$

that have the unique solution

$$\hat{v} = v_3 = g_3 s_4^4.$$

Thus, the reduced model consists of a single reaction $4S_2 + 2S_4 \to 6S_3 + 5S_5$ with $\hat{g}_1 = g_3 S_4^2 / S_2^4$. There is no analogous Kron reduction, because a Kron reduction that prunes species 1 and reactions 1 and 2 would create a direct connection from species 2 to species 3, which is forbidden.

#### S4.2 Comparison of QE and Kron reduction

The QE reduction method consists in pruning a number of fast reactions and pooling species into linear combinations that are conserved by these fast reactions. The number of pools is smaller than the number of pooled species, therefore QE reduces not only the number of reactions but also the number of species.

This method is formalized as follows. Let $R_I$ be a set of fast reactions to prune, and let $I$ denote the set of species that are either produced or consumed by reactions $R_I$. The remaining species $T$ are not affected by $R_I$. Define the set of terminal (slow) reactions as $R_T = \mathcal{V}_r \setminus R_I$. With these definitions, the stoichiometric matrix of the CRN has the following block decomposition:

$$\boldsymbol{S} = \begin{bmatrix} \boldsymbol{S}_I & \boldsymbol{S}_T \\ \mathbb{O} & \boldsymbol{S}_E \end{bmatrix},$$

where $\boldsymbol{S}_T$, $\boldsymbol{S}_E$ contains the stoichiometries of reactions $R_T$ on species $I$ and $T$. Let $\boldsymbol{c} = [c_1, \ldots, c_{n_I}] \in \mathbb{R}^{n_I}$ be a vector of concentrations of species $I$. $\boldsymbol{c}$ is an *approximate conservation law* of $R_I$ if $\boldsymbol{c}\boldsymbol{S}_I = 0$ and $\boldsymbol{c}\boldsymbol{S}_T \neq 0$ [10]. We use the approximate conservation laws to define species pools,

$$\hat{s} = \sum_{i \in I} c_i s_i. \tag{11}$$

A species pool corresponds to a vector $\boldsymbol{c}$ satisfying the following conditions:

*i)* $\boldsymbol{c}\boldsymbol{S}_I = 0$.
*ii)* $\boldsymbol{c}\boldsymbol{S}_T \neq 0$.
*iii)* $c_i \geq 0$ for all $i$, $1 \leq i \leq n_I$.
*iv)* $\boldsymbol{c}$ is minimal, in the sense that if $\boldsymbol{c} = \boldsymbol{c}' + \boldsymbol{c}''$ where $\boldsymbol{c}'$ and $\boldsymbol{c}''$ both satisfy conditions (i)-(iii) and at least one has more zero components than $\boldsymbol{c}$ then $\boldsymbol{c}' = 0$ or $\boldsymbol{c}'' = 0$.

The QE equations are balance equations written for species in $I$, but using only reactions from $R_I$. The rates of the other, slower reactions, are neglected. Namely,

$$\boldsymbol{S}_I \boldsymbol{v}_I = 0, \tag{12}$$

where $\boldsymbol{v}_I$ is the rate vector of reactions $R_I$. Because of the condition i), the matrix $\boldsymbol{S}_I$ is not full rank and Eq. (12) are linearly dependent. However, the rank deficiency is compensated by supplementing them with the conservation equations, which define the pools:

$$\hat{s}_j = \boldsymbol{c}_j \boldsymbol{s}_I,\ 1 \leq j \leq n_C, \tag{13}$$

where $\hat{s}_j$ are the new pool variables $\boldsymbol{s}_I$ is the column vector of concentrations of species $I$, and $n_C$ is the number of independent pools (corresponding to solutions of (i)-(iv)).

To reduce the model, one prunes the reactions $R_I$ and pools the species from $I$ into combinations defined by Eq.(11). The reduced model contains only the terminal reactions $R_T$. The stoichiometries of these reactions are

$$\boldsymbol{c}_j \boldsymbol{S}_T,\ 1 \leq j \leq n_C, \tag{14}$$

on the pools and follow the entries of $\boldsymbol{S}_T$ on species $T$. In summary, the QE reduction algorithm is as follows

**Algorithm S2** QEReduction

**Require:** A CRN given by $\boldsymbol{S}$ matrix and reaction rate vector $\boldsymbol{v}(\boldsymbol{s})$. A subset of reactions $R_I$ to prune.

**Ensure:** A reduced CRN with less reactions and less species, given by a smaller $\hat{\boldsymbol{S}}$ matrix and reaction rate vector $\hat{\boldsymbol{v}}(\hat{\boldsymbol{s}})$.

1: Compute the sets $I$, $T$, $R_T$, and the matrices $\boldsymbol{S}_I$, $\boldsymbol{S}_T$, $\boldsymbol{S}_E$.
2: Eliminate the reactions $R_I$.
3: Compute $\boldsymbol{c}_j$, $1 \leq j \leq n_C \leq n_I$ satisfying i)-iv).
4: Solve (12), (13) for $\boldsymbol{s}_I$.
5: Use solution of (12), (13) obtained at **3.** to express $\boldsymbol{v}_T$ as functions of $\boldsymbol{s}_T$ and $\hat{s}_j$, $1 \leq j \leq n_C$.
6: Define $\hat{\boldsymbol{S}} = \begin{bmatrix} \boldsymbol{c}_1 \boldsymbol{S_T} \\ \vdots \\ \boldsymbol{c}_{n_C} \boldsymbol{S_T} \\ \boldsymbol{S_E} \end{bmatrix}$, $\hat{\boldsymbol{v}} = \boldsymbol{v}_T$, $\hat{\boldsymbol{s}} = \begin{bmatrix} \hat{s}_1 \\ \vdots \\ \hat{s}_{n_C} \\ \boldsymbol{s}_T \end{bmatrix}$

To illustrate these concepts let us consider again the example of the Section 1. Suppose that the fast reactions are $R_I = \{1, 2\}$. In this case the species consumed or produced by these reactions are $I = \{1, 2\}$. The different stoichiometric matrices are:

$$\boldsymbol{S}_I = \begin{bmatrix} -1 & 1 \\ 1 & -1 \end{bmatrix}, \boldsymbol{S}_T = \begin{bmatrix} 0 \\ -1 \end{bmatrix}, \boldsymbol{S}_E = \begin{bmatrix} -1 \\ 1 \end{bmatrix}.$$

$\boldsymbol{c} = [1, 1]$ is an approximate conservation law as it satisfies (i) and (ii). It is also the unique solution of (i)-(iv). In the reduced model the species $I$ are replaced by the single pool variable $\hat{s}_1 = s_1 + s_2$. The reduced model has thus three species variables $\hat{s}_1$, $s_3$, and $s_4$.

The reactions $R_I$ are pruned. The remaining reaction, reaction 3, has stoichiometry $\boldsymbol{c}\boldsymbol{S}_T = -1$ with respect to the pool, and follows the entries of $\boldsymbol{S}_E$ for species 3 and 4. Therefore, the reduced stoichiometric matrix is

$$S_{red} = \begin{bmatrix} -1 \\ -1 \\ 1 \end{bmatrix}.$$

The reaction rate of the remaining reaction, is $v_3$, the same as in the original model. However, it has to be expressed in the new variables. Using QE equation $k_1 s_1 = k_2 s_2$ and the pool definition $\hat{s}_1 = s_1 + s_2$ we get

$$s_1 = \frac{k_2}{k_1 + k_2}\hat{s}_1, \; s_2 = \frac{k_1}{k_1 + k_2}\hat{s}_1.$$

Expressed in the new variables, the reaction rate is

$$v_3 = k_3 s_2 s_3 = \frac{k_1 k_3}{k_1 + k_2}\hat{s}_1 s_3$$

The reduced model reads $\hat{S}_1 + S_3 \to S_4$ with $\hat{g}_1 = \frac{k_1 k_3}{k_1 + k_2}$. This QE approximation cannot be compared to a Kron reduction, because the Kron reduction does not redefine species.

### Acronyms

*1DG* 1-Deoxyosone.
*3DG* 3-Deoxyosone.

*AA* Acetic Acid.
*AMPAR* Activity-dependent Movement of a Glutamate Receptor.

*CDC* Cell Division Cycle.
*CRN* Chemical Reaction Network.

*DFG* N-(1-deoxy-D-fructose-1-yl)-glycine.

*GC* Graph of Complexes.

*HSP* Hedgehog Signaling Pathway.

*MAK* Mass-Action Kinetics.

*ODE* Ordinary Differential Equation.

*pH* Potential of Hydrogen.

*QE* Quasi-Equilibrium.
*QSS* Quasi-Steady-State.

*SR-graph* Species-Reaction Graph.
